# Combinatorial patterns of graded RhoA activation and uniform F-actin depletion promote tissue curvature

**DOI:** 10.1101/2020.04.15.043893

**Authors:** Marlis Denk-Lobnig, Natalie C Heer, Adam C Martin

## Abstract

During development, gene expression regulates cell mechanics and shape to sculpt tissues. Epithelial folding proceeds through distinct cell shape changes that occur in different regions of a tissue. Here, using quantitative imaging in *Drosophila melanogaster*, we investigate how patterned cell shape changes promote tissue bending during early embryogenesis. We find that the transcription factors Twist and Snail combinatorially regulate a unique multicellular pattern of junctional F-actin density, which corresponds to whether cells apically constrict, stretch, or maintain their shape. Part of this pattern is a gradient in junctional F-actin and apical myosin-2, and the width of this gradient regulates tissue curvature. The actomyosin gradient results from a gradient in RhoA activation that is refined by a balance between RhoGEF2 and the RhoGAP C-GAP. Thus, cell behavior in the ventral furrow is choreographed by the interplay of distinct gene expression patterns and this coordination regulates tissue shape.

## Introduction

During development, the three-dimensional shape of a complex organism is generated, in part, by gene expression patterns that are encoded by a one-dimensional sequence of nucleotides in the genome. Patterns of gene expression and resulting signaling processes overlap and interact in space and time to define each cell’s function. For example, morphogen gradients encode positional information for specific cell fates (Rogers and Schier, 2011; Wolpert, 1969). For tissues to obtain their final and functional state, cell fates, shapes, and mechanics all need to be positionally specified. Cell fate and mechanical patterns need not be identical, as mechanical properties are often patterned within cells of the same type (Mongera et al., 2018; Sui et al., 2018; Sumigray et al., 2018). Each tissue shape change requires coordinated changes in cell shape and position across the tissue, which have to be tailored to the tissue’s morphological and functional requirements while being robust and reproducible between individual embryos/organisms (Chanet et al., 2017; Hong et al., 2016; von Dassow and Davidson, 2009).

Mesoderm invagination in the early *Drosophila melanogaster* embryo involves folding an epithelial sheet and is an established model system for gene expression patterning and morphogenesis (Leptin, 2005). Mesoderm invagination requires apical constriction, a cell shape change driven by actomyosin constriction that converts columnar epithelial cells to a wedge shape, which promotes tissue bending (Leptin and Grunewald, 1990; Sweeton et al., 1991). Importantly, apical constriction is coordinated across the presumptive mesoderm; there is a spatial, ventral-lateral gradient of apical non-muscle Myosin-2 (myosin) and apical constriction that extends ∼ 5 - 6 cell rows from the ventral midline (Heer et al., 2017; Lim et al., 2017; Oda and Tsukita, 2001; Spahn and Reuter, 2013) (Fig. 1 A). Beyond this gradient, apical myosin levels reach a baseline low level, and 2 - 4 cell rows (rows ∼ 7 – 9) stretch their apical surface and bend towards the forming furrow (Heer et al., 2017; Leptin and Grunewald, 1990; Sweeton et al., 1991). In contrast, more lateral cells, which are part of the neighboring ectoderm and also have baseline apical myosin levels, maintain an almost constant apical area throughout the folding process (Rauzi et al., 2015). We investigated how this tissue-wide pattern of cell shapes is established and how it impacts the resulting tissue shape.

**Figure 1.**
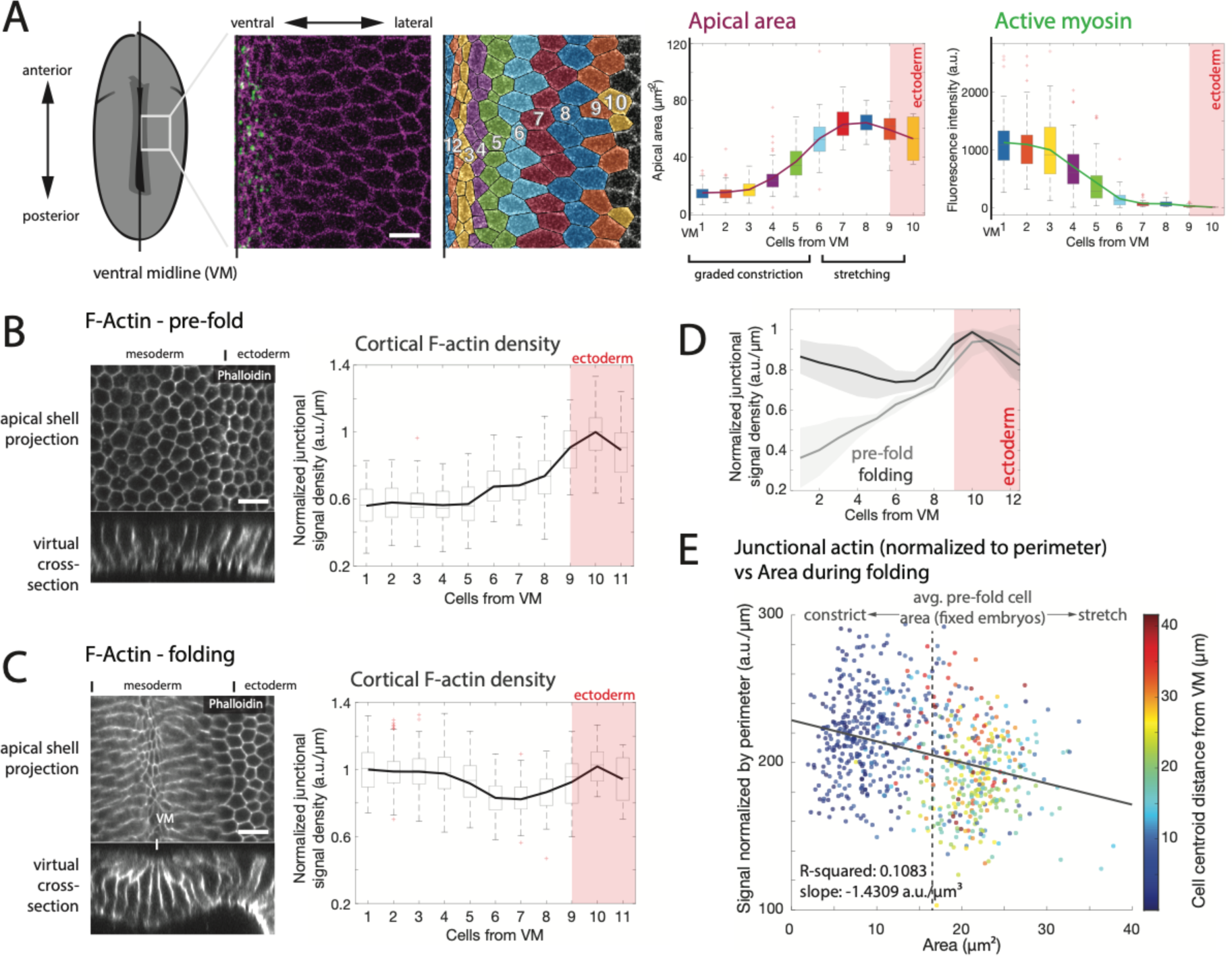
Tissue-wide mesodermal F-actin distribution is distinct from Myosin. **(A)** Myosin gradient extends 6 cells from the ventral midline (VM) and remains low. Images are apical surface view of embryo labeled with sqh::GFP (Myosin) and Gap43::mCherry (membrane) and segmented example of embryo with cell rows highlighted in different colors. Scale bar = 10 µm. Colors in image correspond to bins in adjacent plots showing myosin intensity and apical area distributions in a representative wild-type embryo. **(B)** Junctional F-actin is depleted in the mesoderm prior to furrow formation. Images are apical shell projection (top) and cross-section (bottom) image of phalloidin stained embryos imaged before furrow formation. Scale bar = 10 µm. Plot shows junctional F-actin density for cells of one embryo binned by position from ventral midline. Data is represented by box-and-whisker plot from representative embryo, where bottom and top sides of the box represent 25^th^ and 75^th^ percentile of cells, respectively. Midline is the median and red ‘+’ are outliers. n = at least 40 cells per bin (median 51.5 cells). **(C)** Junctional F-actin accumulates in a ventral-lateral gradient during furrow formation. Images and plots are analogous to (B). n = at least 15 cells per bin (median 73 cells). **(D)** Average F-actin density trace for 3 embryos corresponding to (B) (pre-furrow) and (C) (furrow), normalized to the cell row with the highest mean. Shaded area depicts 1 standard deviation in each direction. **(E)** Apical area is anti-correlated with cortical F-actin levels. Quantification of cortical F-actin density per cell (cortical levels normalized by perimeter) for the embryo shown in (C) was plotted as a function of apical area. Color of data points represents physical distance from ventral midline in µm. Average pre-fold cell area for fixed embryos (∼16 µm^2^ due to shrinkage during the fixation process) is indicated with dotted grey line.

Mesoderm cell shape change and cell fate are initially driven by the transcription factors Dorsal, Twist, and Snail (Boulay et al., 1987; Furlong et al., 2001; Leptin, 1991; Thisse et al., 1988), which exhibit distinct expression patterns. Nuclear Dorsal is present in a ventral-dorsal gradient that narrows over time (Rahimi et al., 2019; Roth et al., 1989; Rushlow et al., 1989; Steward, 1989; Steward et al., 1988). Measurements of the transcription dynamics of Twist target genes, *T48* and *fog*, demonstrate that downstream Twist targets are expressed first at the ventral midline and subsequently expand laterally (Lim et al., 2017). This temporal progression of gene expression results in the graded accumulation of T48 transcripts and protein along the ventral-lateral axis (Heer et al., 2017; Lim et al., 2017; Rahimi et al., 2019). Snail can both activate and repress gene expression (Rembold et al., 2014). One gene that is activated by Snail is the G-protein coupled receptor (GPCR) Mist (Manning et al., 2013). In contrast to graded Twist activity, Snail activity as assessed by Mist mRNA expression is uniform across the mesoderm (Lim et al., 2017). Therefore, Twist and Snail target genes appear to have distinct patterns of expression.

The product of the Twist target gene *fog* activates the Mist GPCR and a uniformly expressed GPCR, Smog (Costa et al., 1994; Kerridge et al., 2016; Manning et al., 2013). This GPCR pathway and T48 expression act via the guanine nucleotide exchange factor (GEF) RhoGEF2 and the small GTPase RhoA to activate myosin contractility (Barrett et al., 1997; Hacker and Perrimon, 1998; Kolsch et al., 2007). Functioning in opposition to RhoGEF2 to shut off RhoA signaling activity is the RhoA GTPase-activating protein (GAP) C-GAP (also called Cumberland-GAP or RhoGAP71E) (Mason et al., 2016). RhoA coordinately activates myosin, via Rho-associated and coiled coil kinase (ROCK), and actin filament (F-actin) assembly, via the formin Diaphanous (Dawes-Hoang et al., 2005; Homem and Peifer, 2008). Myosin activation occurs in a gradient that is narrower than the gradient of T48 protein accumulation (Heer et al., 2017), but it is not known whether RhoA activation follows a similar pattern to T48 or whether RhoA signaling is refined by its regulators. F-actin is apically enriched in ventral cells in a manner that depends on RhoA signaling (Fox and Peifer, 2007). We asked whether there is a multicellular pattern to F-actin across the mesoderm and how this is controlled transcriptionally and via RhoA signaling.

Here we show a tissue-level pattern of junctional F-actin density that is distinct from myosin activation and Twist activity. This pattern of junctional F-actin results from the combination of Snail-dependent depletion and Twist-dependent accumulation, which is tuned by RhoA activity level. We show that the width of the actomyosin gradient regulates tissue curvature, as well as lumen size in the mesodermal tube structure generated by this fold. Our results show how combinatorial patterning of two transcriptional programs creates distinct zones of cytoskeletal protein accumulation across the mesoderm and that this patterning promotes proper tissue curvature.

## Results

### Junctional F-actin exhibits a distinct tissue pattern from apical myosin during mesoderm invagination

To determine how the ventral-dorsal pattern of F-actin changes during mesoderm invagination, we labeled F-actin in fixed or live embryos with phalloidin or Utrophin::GFP, respectively, and measured F-actin in cell bins at defined positions from the ventral midline. We chose to focus our analysis on junctional F-actin density, because cortical F-actin density is known to affect cell mechanics (Salbreux et al., 2012; Stricker et al., 2010) and we observed clear variation of junctional F-actin across the mesoderm and neighboring ectoderm. Before ventral furrow formation onset, mesodermal junctional F-actin density (and integrated F-actin intensity) drops relative to ectoderm cells (Fig. 1 B and D and Supplemental Fig. 1 A - C), consistent with what has been previously observed (Jodoin et al., 2015).

During ventral furrow formation, junctional F-actin density rose around the ventral midline and exhibited a gradient that extended to cell 6, similar to apical myosin (Fig. 1 C, D, and Supplemental Fig. 1 A - C). F-actin density remained lower in cell rows ∼ 6 - 8 of the mesoderm, forming a zone of junctional F-actin depletion relative to the neighboring ectoderm. This contrasts with apical myosin activation, which decreases to a baseline level in marginal mesoderm cells that is similar to the neighboring ectoderm (Fig. 1 A) (Heer et al., 2017; Lim et al., 2017; Spahn and Reuter, 2013). The observed pattern was also present in live embryos expressing the actin-binding domain of Utrophin fused to green fluorescent protein (Utrophin::GFP) or mCherry (Utrophin::mCherry), but not with a general membrane marker, Gap43-mCherry, indicating that this measurement is not a fixation artifact or due to changes in plasma membrane structure, such as stretching (Supplemental Fig. 1 D). Thus, junctional F-actin levels exhibit a distinct pattern from that of apical myosin activation.

The observed tissue-wide pattern of F-actin density matched the pattern of apical cell area constriction and stretching (Fig. 1 A, C; Supplemental Fig. 1A). To quantify the strength of this relationship, we correlated junctional cell F-actin density with apical cell area in both mesoderm and adjacent ectoderm cells (Fig. 1 E). We found an anti-correlation between junctional F-actin density and apical area (R-squared: 0.1083), in part, because marginal mesoderm cells (green/yellow points) had low F-actin and were stretched and adjacent ectoderm cells (red) had intermediate F-actin and did not stretch or constrict. Because F-actin density and turnover influence the ability to dissipate stress (Clement et al., 2017; Salbreux et al., 2012; Stricker et al., 2010), our result suggests that lower F-actin levels allow lateral mesoderm cells to stretch, resulting in an inverted cell morphology for cells at the edge of the mesoderm compared to apically constricted cells at the ventral midline.

### Snail and Twist regulate distinct components of the F-actin pattern

To determine how this tissue-wide F-actin pattern is established, we tested how the transcription factors *snail* and *twist* affect junctional F-actin density. Snail activity in the mesoderm, as measured by *mist* transcription, is uniform (Lim et al., 2017) and we found that the Snail boundary co-localizes precisely with the F-actin depletion boundary (Fig. 2 A). Furthermore, mesodermal cell junctional F-actin depletion requires *snail*. To illustrate and quantify junctional F-actin around the curved embryo, we created a reslice at a constant height from the apical surface (Supplemental Fig. 2). Unlike control (heterozygous) embryos, *snail* homozygous mutant embryos expressing fluorescently tagged Utrophin do not exhibit patterned junctional F-actin levels in the ventral region, but rather uniform intensity throughout the region, without a sharp boundary (Fig. 2 B and C). Because *snail* mutant embryos were imaged live, we could identify the mesoderm/ectoderm boundary based on subsequent germband extension movements in the ectoderm and premature cell divisions that occurred in the uninternalized mesoderm, which is mitotic domain 10 (Foe, 1989; Grosshans and Wieschaus, 2000). We conclude that Snail decreases junctional F-actin density in the mesoderm.

**Figure 2.**
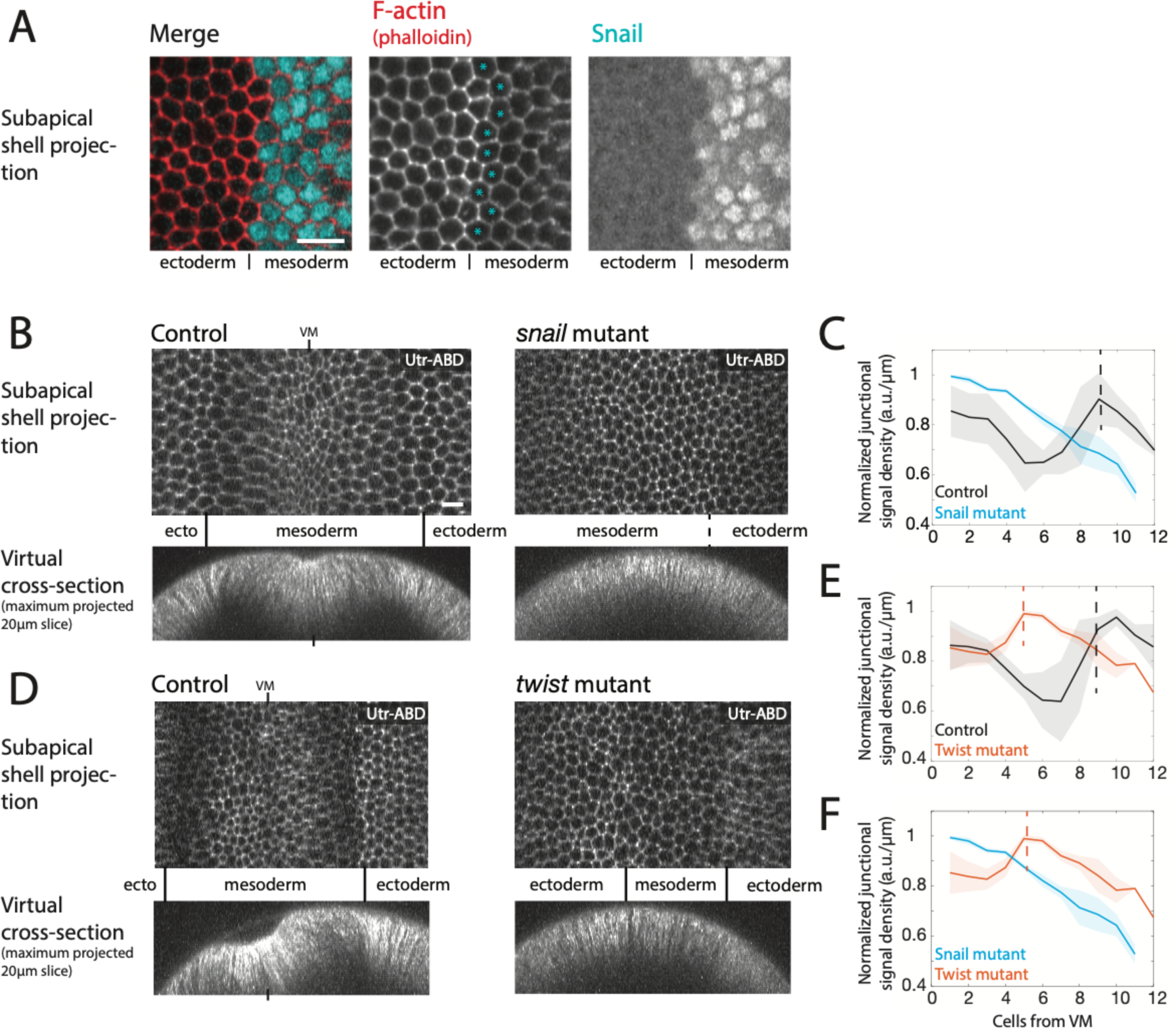
Snail and Twist regulate distinct features of the tissue-wide F-actin pattern. **(A)** Snail expression boundary corresponds to F-actin depletion boundary. Images are from phalloidin-stained embryo co-stained with anti-Snail. Cyan stars in F-actin image designate Snail-positive cells bordering the ectoderm. Scale bar = 10 µm. **(B)** The *snail* mutant disrupts mesodermal F-actin depletion. Images are subapical shell-projections from representative, live homozygous *snail* mutant and normal sibling embryo expressing Utrophin::mCherry (F-actin). The *snail* mutant shows a lack of F-actin patterning. Scale bar = 10 µm. **(C)** Quantification of junctional F-actin density by cell row from 3 *snail* mutant and 3 normal sibling embryos (Mean and standard deviation shown). All *snail* mutants lack F-actin patterning. All traces were normalized to their highest-mean cell bin before averaging. Dotted black line designates approximate mesoderm-ectoderm boundary in control embryos. **(D)** The *twist* mutant exhibits F-actin depletion, but lacks F-actin elevation around midline. Images are subapical shell-projections from 3 representative, live homozygous *twist* mutant and 3 normal sibling embryos expressing Utrophin::mCherry (F-actin). Scale bar = 20 µm. **(E)** Quantification of junctional F-actin density from 3 *twist* mutant and 3 normal sibling embryos (Mean and standard deviation shown). All *twist* mutants exhibit F-actin depletion, but not the increase around the midline. All traces were normalized to their highest-mean cell bin before averaging. Dotted lines mark respective transitions from low-F-actin to high F-actin regions of the tissue. **(F)** Average F-actin density and standard deviations overlaid for 3 *twist* and 3 *snail* mutant embryos. *twist* mutants display ventral F-actin depletion but *snail* mutants do not.

In contrast to Snail, the expression of Twist transcriptional targets is graded, with Twist target expression initiating first along the ventral midline and then expanding laterally (Lim et al., 2017; Rahimi et al., 2019). Twist also regulates *snail* expression; *twist* mutants reduce *snail* expression width (Leptin, 1991). To determine *twist*’s requirement for the tissue-wide pattern of junctional F-actin, we examined F-actin density in a *twist* null mutant that has been shown to disrupt myosin stabilization, but still exhibits transient actomyosin contractions (Martin et al., 2009). In contrast to *snail* mutants, *twist* mutant embryos exhibit lower junctional F-actin density in the mesoderm and a clear boundary with the ectoderm (Fig. 2 D, E, and F). However, the zone of low junctional F-actin is decreased to half the normal width, consistent with narrowed *snail* expression depleting F-actin in a narrower mesoderm. Graded F-actin accumulation around the ventral midline was absent in *twist* mutants, suggesting that higher F-actin density at the midline depends on the Twist pathway, which includes RhoA activation (Dawes-Hoang et al., 2005; Fox and Peifer, 2007; Kolsch et al., 2007; Mason et al., 2013). Therefore, mesodermal control of junctional F-actin by Twist and Snail is comprised of two nested layers: 1) prior to ventral furrow formation, uniform Snail activity lowers F-actin density across the mesoderm, and 2) during ventral furrow formation there is a Twist-dependent increase in F-actin density that, similarly to apical myosin, creates a ventral-lateral gradient, but also creates a ‘valley’ in junctional F-actin density at the margin of the mesoderm.

### Neither graded myosin activation nor F-actin depletion depend on intercellular mechanical connections or cell shape change

Our data suggested that Snail and Twist promote uniform mesodermal F-actin depletion prior to apical constriction and subsequent graded actomyosin accumulation, respectively. In the ventral furrow and the related process of *Drosophila* posterior midgut formation, it has been proposed that mechanical feedback between constricting cells is required to induce myosin accumulation, which may contribute to a gradient or wave in contractility (Bailles et al., 2019; Mitrossilis et al., 2017). Therefore, it was important to determine whether transcriptional activity is directly shaping these patterns or whether mechanical induction of myosin contributes as well.

To determine the contribution of intercellular mechanical connections to the multicellular patterns of myosin activation and junctional F-actin density, we examined embryos in which intercellular cytoskeletal coupling was disrupted. We did this by depleting the adherens junction protein α-catenin by RNA interference (α-catenin-RNAi), which uncouples the cytoskeletal meshwork of cells from the junctions and, thus, mechanical connections between cells (Fig. 3 A, Supplemental Movie S1, 2) (Fernandez-Gonzalez and Zallen, 2011; Martin et al., 2010; Yevick et al., 2019). In α-catenin-RNAi embryos, ventral cell apical area remains at pre-gastrulation levels (about 40 μm^2^), but myosin contracts into spot-like structures (Fig. 3 A, B, E) (Martin et al., 2010). Apical myosin quantification showed that, despite the lack of mechanical coupling, myosin reproducibly accumulates in a gradient around the ventral midline that is similar to the wild-type gradient (Fig. 3 C, E). Therefore, graded myosin activation across the tissue does not depend on force transmission between cells.

**Figure 3.**
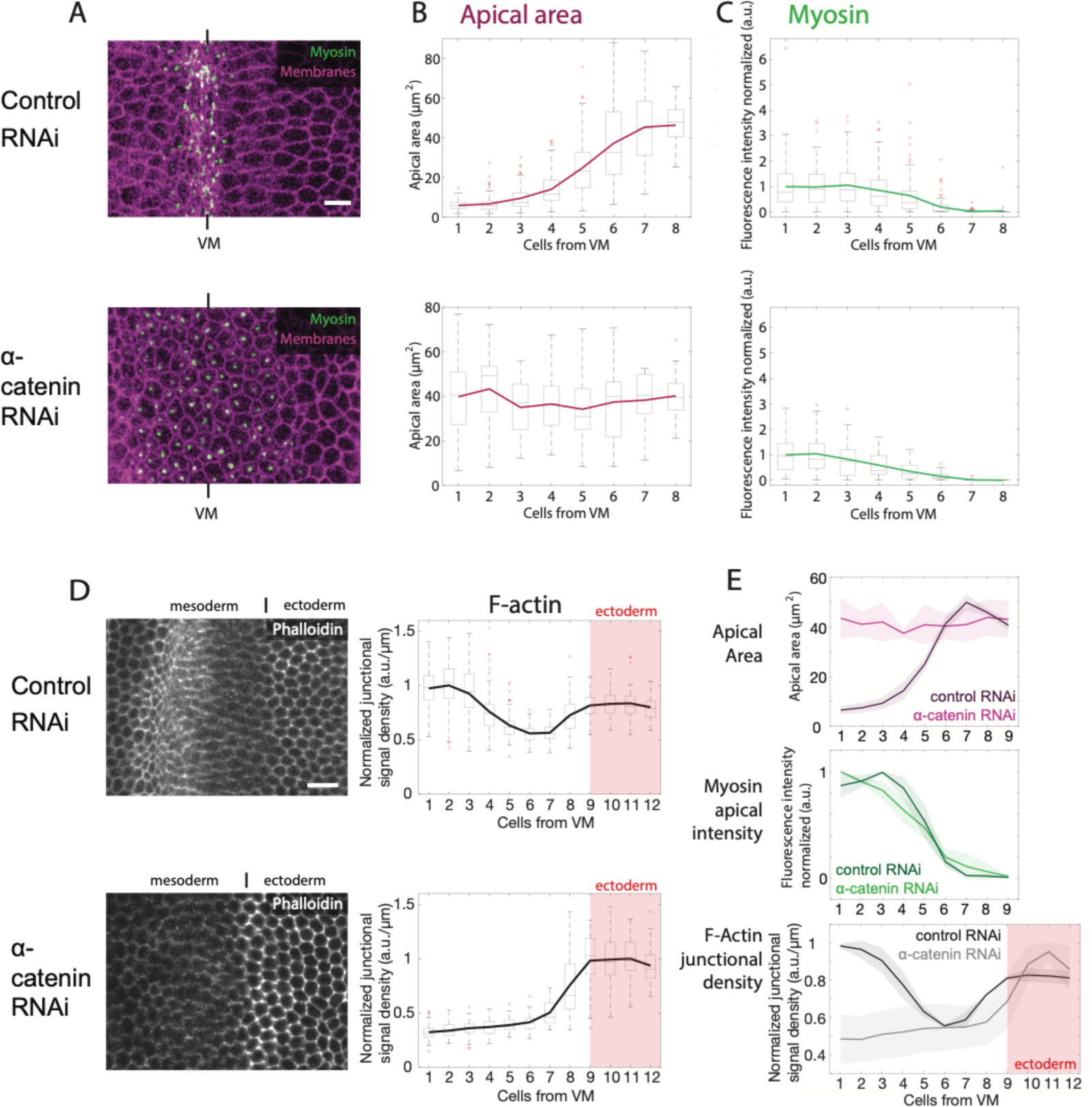
Myosin gradient and uniform F-actin depletion do not require intercellular coupling. **(A)** Images (apical shell projections) of control (Rh3-RNAi) and α-catenin-RNAi embryos expressing sqh::GFP (Myosin, green) and Gap43::mCherry (Membranes, magenta). Scale bar = 10 µm. **(B)** and **(C)** Quantification of apical area (magenta, **B**) and normalized apical, active myosin (green, **C**) as a function of distance from ventral midline. Data is represented by box-and-whisker plots where each bin is a cell row at a given distance from the ventral midline (n = at least 41 (control) or 44 (α-catenin-RNAi) cells per row; median 72 (control) or 107 (α-catenin-RNAi) cells). Bottom and top sides of the box represent 25^th^ and 75^th^ percentile, respectively. Midline is the median and red points are outliers. **(D)** Images are phalloidin-stained control (Rh3-RNAi, top) and α-catenin-RNAi (bottom) embryos focused on the mesoderm-ectoderm boundary. Scale bar = 10 µm. Plots are normalized junctional F-actin density in rows of cells along the ventral-lateral axis (n= at least 52 (control) or 8 (α-catenin-RNAi) cells per row; median 78 (control) or 40.5 (α-catenin-RNAi) cells). Data representation is same as (B) and (C). **(E)** Mean (+ standard deviation) traces for several control and α-catenin-RNAi embryos. Top: Average apical area behavior from 4 different embryos, per condition. Mid: Average apical myosin intensity behavior from 4 different embryos, respectively. Bottom: Average junctional actin density from 3 different embryos, respectively.

Adherens junctions are a known target of Snail in the mesoderm (Chanet and Schweisguth, 2012; Dawes-Hoang et al., 2005; Kolsch et al., 2007) and E-cadherin exhibits a similar tissue-level pattern to F-actin (Supplemental Fig. 3A). Therefore, we asked whether mesodermal F-actin depletion depended on intact adherens junctions. Similar to wild-type embryos, junctional F-actin density in α-catenin-RNAi embryos is low across the mesoderm with a sharp boundary to the ectoderm (Fig. 3 D, E). This observation suggests that the Snail-mediated F-actin density reduction does not depend on intact junctions. Furthermore, low F-actin density in lateral mesoderm cells is maintained despite a lack of apical constriction and stretching of lateral mesoderm cells, confirming that reduced F-actin density in these cells is not due to cell stretching. Interestingly, junctional F-actin levels around the midline does not appear to increase the same in α-catenin-RNAi embryos as in wild-type embryos during constriction. The lack of increased F-actin density may be due to disrupted intercellular junctions in the α-catenin-RNAi embryos, but it is also possible that the increased junctional F-actin density at the midline in wild-type/control embryos is due to apical constriction. Medio-apical F-actin foci, which co-localize with myosin, appear in α-catenin-RNAi embryos, and it is possible that elevated junctional F-actin is drawn in to these structures (Supplemental Fig. 3 B). These results indicate that F-actin depletion in the mesoderm does not require intact adherens junctions or mechanical coupling of cells, which is consistent with transcriptional regulation causing the observed pattern.

### RhoA activation occurs in a gradient

We next investigated further what is responsible for the gradient in apical myosin and junctional F-actin. Fog and T48 transcripts accumulate in a gradient (Lim et al., 2017; Rahimi et al., 2019), but the width of T48 protein accumulation is wider than the gradient in apical myosin (Heer et al., 2017). Because Fog and T48 function upstream of RhoA, we examined fluorescently tagged versions of RhoA’s activator RhoGEF2 (under an endogenous promoter), the Anillin Rho-binding domain (an active RhoA sensor), and the RhoA effector ROCK (Mason et al., 2016; Munjal et al., 2015; Simoes Sde et al., 2010) (Fig. 4 A). Each of these fluorescent protein markers became apically enriched in ventral cells during ventral furrow formation, consistent with previous studies (Fig. 4 B) (Kolsch et al., 2007; Mason et al., 2013; Mason et al., 2016).

**Figure 4.**
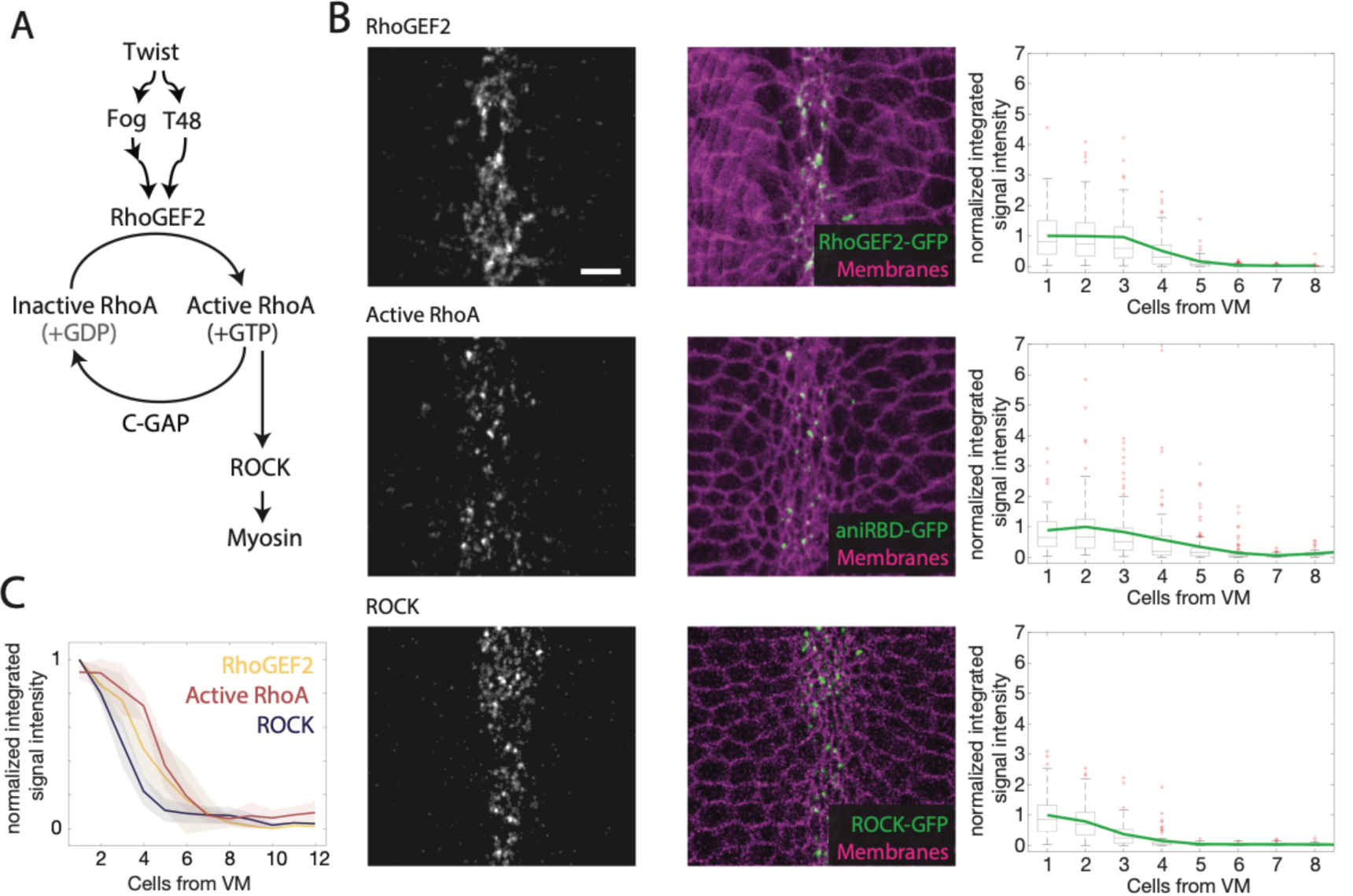
RhoA activation occurs in a gradient. **(A)** Simplified diagram of signaling downstream of Twist, focused on RhoA regulation. **(B) Top**: Image of RhoGEF2::GFP, **Middle:** Image of Anillin Rho-binding domain::GFP (Active RhoA), **Bottom:** Image of Rok::GFP, and Gap43::mCherry (membranes). Ventral midline in center. Scale bar = 10 µm. Plots are normalized apical RhoGEF2::GFP intensity (**top**), Anillin Rho-binding domain::GFP intensity (**mid**), and Rok::GFP intensity (**bottom**) as a function of distance from the ventral midline for one representative embryo, respectively. Data is represented by box-and-whisker plots where each bin is a cell row at a given distance from the ventral midline. Bottom and top sides of the box represent 25^th^ and 75^th^ percentile, respectively. Midline is the median and red points are outliers. At least 32 cells (rhoGEF2, median 85 cells), 58 cells (Active RhoA, median 76.5 cells), or 51 cells (Rock, median 68.5 cells) were analyzed for each cell row. Brightness and contrast were adjusted individually to best display the intensity range for each marker. **(C)** Average (+ standard deviation) fluorescent signal across cell rows for 5 (RhoGEF2::GFP, Rok::GFP) or 4 (Anillin Rho-binding domain::GFP) embryos, respectively. Traces are normalized to highest-mean cell row, respectively.

Quantification of apical fluorescence by cell row revealed that all three markers for RhoA pathway activation were graded along the ventral-lateral axis, exhibiting strong fluorescence at the ventral midline and gradually decreasing to baseline after ∼ 6 cells (Fig. 4 B and C). In contrast, endogenously tagged C-GAP-GFP was largely cytoplasmic and appeared uniform across the ventral domain during folding (Supplemental Fig. 4 A). Therefore, there is a ventral-lateral gradient of RhoA activation in the mesoderm. We next tested how the RhoA gradient affects graded apical myosin and junctional F-actin.

### RhoGEF and GAP modulate actomyosin gradient width

The gradient in accumulated T48 transcripts and protein extends beyond the 5-6 cell rows from the ventral midline where we detect RhoA and myosin activation (Heer et al., 2017; Lim et al., 2017; Rahimi et al., 2019). Therefore, we hypothesized that the signaling network downstream of T48 further shapes the contractile gradient. Mesodermal RhoA signaling involves a signaling circuit, which includes an activator/inhibitor pair (Fig. 4 A). The activator RhoGEF2 is required for high levels of apical myosin accumulation (Dawes-Hoang et al., 2005; Nikolaidou and Barrett, 2004). Severely depleting the inhibitor, C-GAP, disrupts proper subcellular myosin localization (Mason et al., 2016). To test the importance of this circuit in regulating the multicellular patterning of the actin cytoskeleton, we either depleted or overexpressed the RhoA regulators C-GAP or RhoGEF2, and determined the effect on actomyosin distribution.

First, we examined apical myosin after elevating RhoA activation by either depleting C-GAP by RNAi (C-GAP-RNAi) or overexpressing RhoGEF2 (RhoGEF2 O/E). These perturbations elevate myosin in mesoderm cells, whereas there is still no noticeable myosin accumulation in the ectoderm at this stage (Fig. 5 A, B). C-GAP-RNAi and RhoGEF2 O/E expand the myosin gradient width, with myosin activation occurring 1 – 2 cell rows farther from the ventral midline than control embryos (Fig. 5 C, D). C-GAP-RNAi also expands the zone of upstream RhoA activation and downstream apical constriction around the ventral midline (Fig. 5 C and Supplemental Fig. 5 A). Similar to C-GAP-RNAi, RhoGEF2 O/E expands the zone of apical constriction, although cells at the ventral midline constrict less than in control embryos (Fig. 5 D). Interestingly, decreasing RhoA activation by RhoGEF2 depletion (RhoGEF2-RNAi) does not narrow the myosin gradient, although the overall levels and uniformity of cellular myosin (and constriction) are decreased (Supplemental Fig. 5 B). Overall, our data show that increasing RhoA activity increases the width of the myosin gradient, with C-GAP playing an integral role in restricting myosin gradient width.

**Figure 5:**
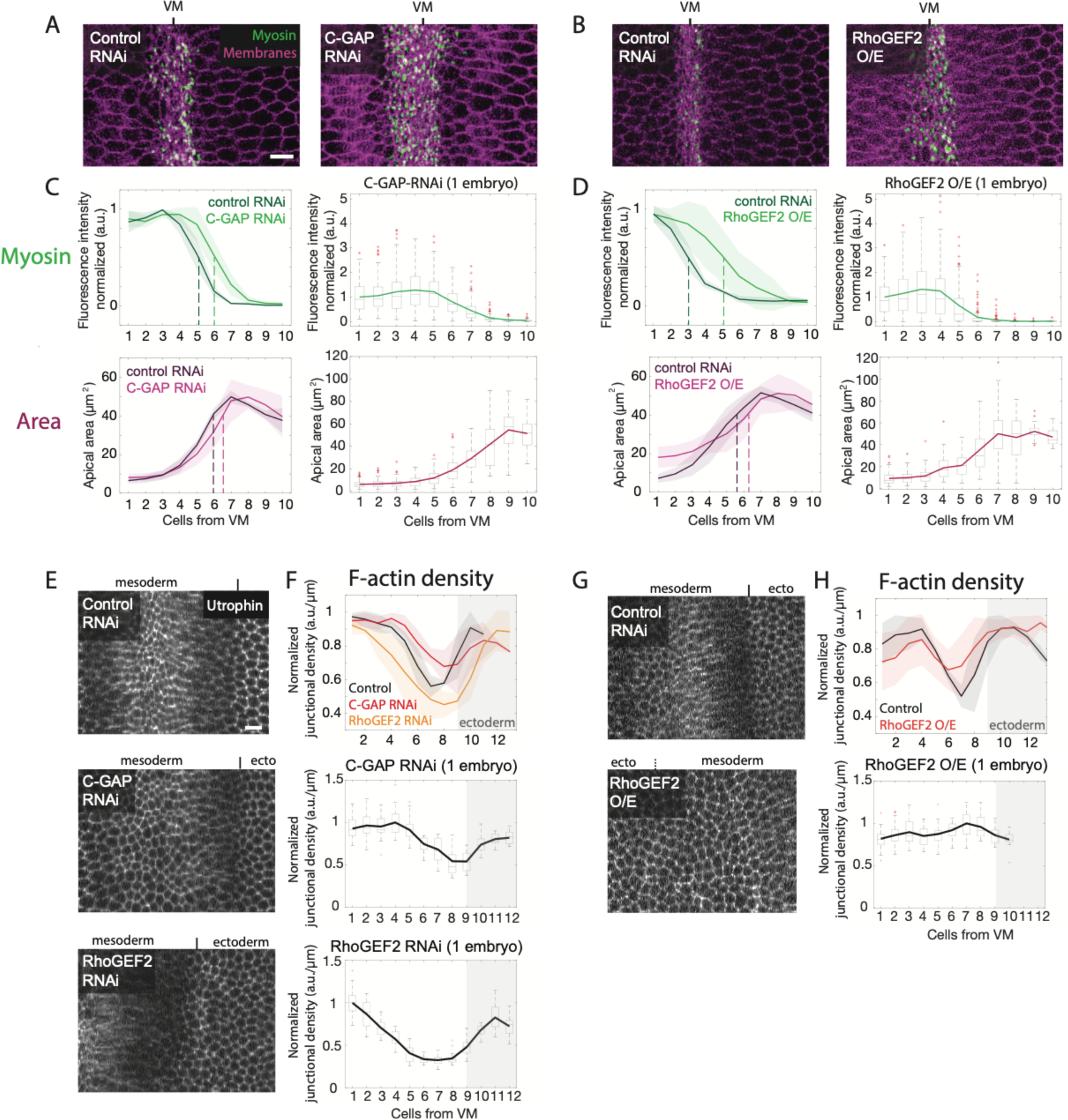
RhoA overactivation widens actomyosin gradient and elevates F-actin at the mesoderm margin. **(A-B)** Images (apical shell projections) of control (Rh3-RNAi) and C-GAP-RNAi embryos **(A)** expressing two copies of sqh::GFP (Myosin, green) and Gap43::mCherry (Membranes, magenta), or control and RhoGEF2 O/E embryos **(B)** expressing one copy of sqh::GFP (Myosin, green) and Gap43::mCherry (Membranes, magenta). Scale bars = 10 µm. **(C-D)** C-GAP-RNAi and RhoGEF2 O/E embryos have a wider half-maximal gradient position. **Left:** Average (+ standard deviation) normalized myosin intensity (top) and apical area (bottom) behavior for **(C)** control (n = 4 embryos) and C-GAP RNAi (n = 5 embryos) or **(D)** control (n = 4 embryos) and RhoGEF2 O/E (n = 7 embryos). Dashed lines indicate half-maximal myosin and unconstricted apical area, respectively. Control embryos for (C) are the same as in Fig. 3 E. **Right:** Quantification of normalized apical, active myosin (green, **top**) and apical area (magenta, **bottom**) as a function of distance from ventral midline for a single C-GAP-RNAi (C) or RhoGEF2 O/E embryo (D). Data is represented by box-and-whisker plots where each bin is a cell row at a given distance from the ventral midline (n= at least 50 cells/row (C) or 23 cells/row (D); median 108.5 cells (C) and 85 cells (D)). Bottom and top sides of the box represent 25^th^ and 75^th^ percentile, respectively. Midline is the median and red points are outliers. **(E** and **G)** Images (subapical shell projection) of control (Rh3-RNAi), C-GAP-RNAi, or RhoGEF2-RNAi embryos (E) or RhoGEF2 O/E embryo (G) expressing Utrophin::GFP. Scale bars = 10 µm. **(F** and **H)** RhoA activity affects junctional F-actin density in cells at the mesoderm margin. **Top:** Average F-actin density behavior (and standard deviation) from different embryos, normalized to mean of highest cell row, for **(F)** control RNAi (n=3), C-GAP-RNAi (n=3), and RhoGEF2-RNAi (n=3) or **(H)** control RNAi (n=2), RhoGEF2 O/E (n=5). **Mid:** Quantification of normalized subapical F-actin density as a function of distance from ventral midline for a single C-GAP-RNAi **(F)** or RhoGEF2 O/E **(H)** embryo (n= at least 9 cells/row (F and H); median 25.5 cells (F) and 21.5 cells (H)). Data representation same as (C) and (D). **Bottom:** Quantification of normalized subapical F-actin density as a function of distance from ventral midline for a single RhoGEF2-RNAi embryo (n= at least 11 cells/row; median 15 cells).

To determine whether RhoA activation is critical for regulation of the junctional F-actin pattern, we examined junctional F-actin density after either elevating or depleting RhoA activity. Elevating RhoA activation, either by C-GAP-RNAi or RhoGEF2 O/E increases junctional F-actin density in marginal mesoderm cells, causing them to be more similar to ectoderm cells (Fig. 5 E-H). In some cases, RhoGEF2 O/E completely eliminates the pattern of F-actin depletion and elevation across the mesoderm (Fig. 5 G, H). Conversely, decreasing RhoA activity by RhoGEF2-RNAi further depletes junctional F-actin in marginal mesoderm cells (Fig. 5 F). Overall, our data suggest that RhoA activity regulates myosin gradient width and also the pattern of junctional F-actin density across the mesoderm.

### Myosin pulses elicit different area responses in midline versus marginal mesoderm cells

Myosin pulses are discrete events in which there is a burst of myosin accumulation and constriction of the cell apex. RhoA activity levels determine the contractile outcome of a myosin pulse in individual cells during ventral furrow formation (Mason et al., 2016). Given that RhoA activity is graded across the furrow (Fig. 4), we examined the behavior of myosin pulses throughout the mesoderm. There are distinct classes of myosin pulses that occur during furrow formation. ‘Ratcheted pulses’ are events in which apical, active myosin persists after a pulse and decreased apical area is sustained (Fig. 6 A) (Xie and Martin, 2015). In contrast, ‘unratcheted pulses’ exhibit myosin dissipation after the pulse and cell relaxation follows constriction. There is a continuum of behaviors from ratcheted to unratcheted, which are associated with high and low RhoA activity, respectively (Mason et al 2016).

**Figure 6:**
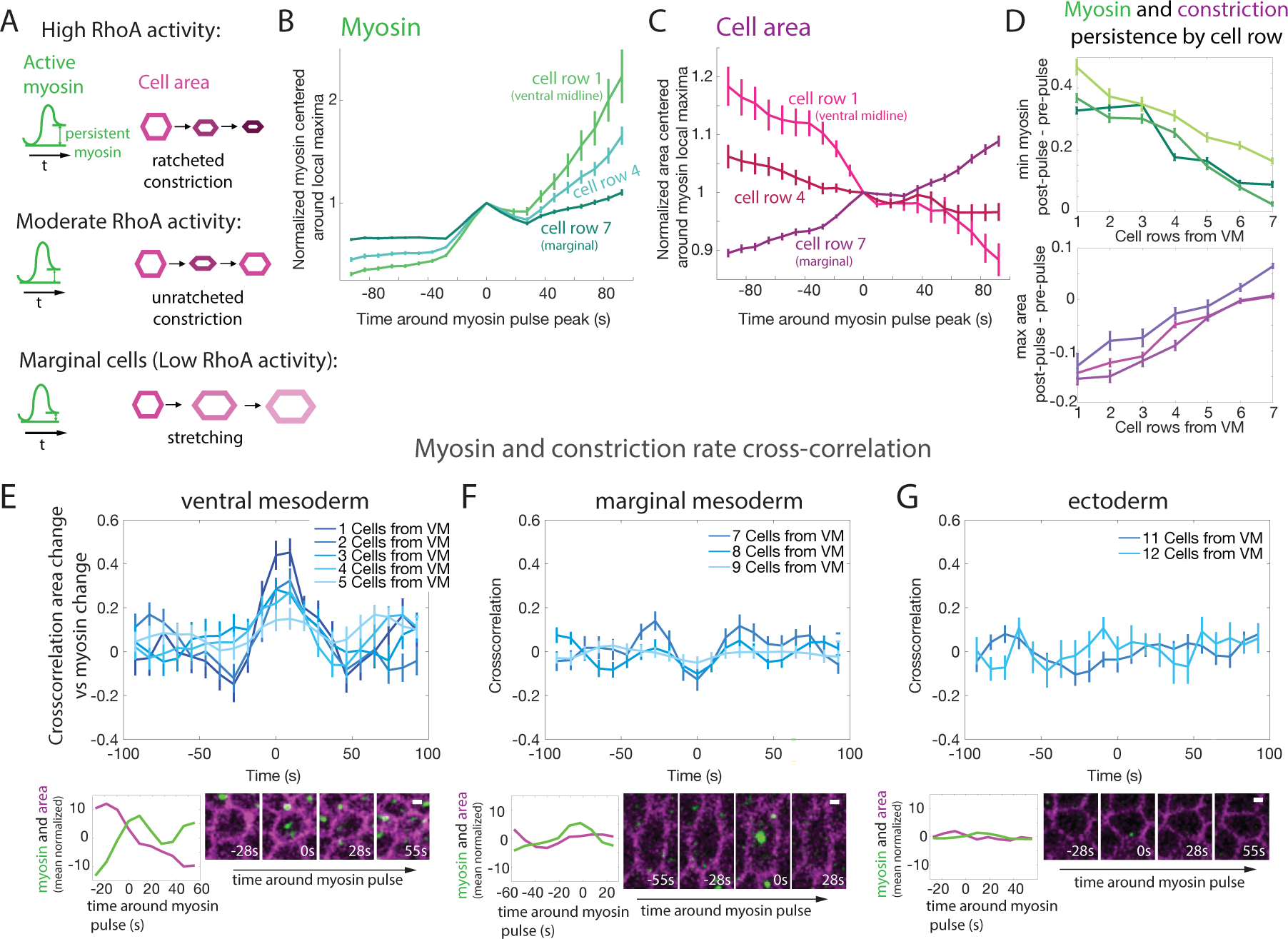
Contractile dynamics vary with distance from ventral midline. **(A)** Diagram categorizing types of dynamic cell behaviors during ventral furrow formation. Top: When RhoA activity is high, cells preferably undergo “ratcheted” pulses after which a higher myosin constriction level is maintained than before the pulse. At moderate RhoA activity, cells preferably undergo reversible pulses in which myosin and constriction levels are similar to before the pulse. **(B-C)** Average and standard error of myosin dynamics **(B)** and apical area dynamics **(C)** around pulses (local maxima in myosin accumulation rate) for cell row 1 at the midline, cell row 4 within the myosin gradient and marginal mesoderm cell row 7 shown for one representative embryo. 106 pulses were analyzed for cell row 1; 192 pulses were analyzed for cell row 4; 335 pulses were analyzed for cell row 7. **(D)** Average and standard error for persistence of myosin (minimum myosin 0-100 s after pulse - minimum myosin 0-100 s before pulse) and area (maximum area 0-100 s after pulse - maximum area 0-100 s before pulse) by bin (distance from the midline) for three embryos. At least 83 pulses were analyzed for each cell row in each embryo. Median 192, 238 and 343 pulses analyzed per cell row for the three embryos, respectively. **(E)-(G)** Top: Cross-correlation of myosin rate and constriction rate averaged by cell bin; split up by ventral mesoderm (cell rows 1-5, **E**), marginal mesoderm (cell rows 7-9, **F**) and ectoderm (cell rows 11 and 12, **G**). At least 21 cells per cell row were analyzed, median 32 cells per cell row. Bottom: Myosin (green) and apical area (magenta) traces (normalized to average) and images of representative individual cells during a myosin pulse, for each region. Scale bars = 2 µm.

To determine whether the ventral-lateral gradient in RhoA patterns myosin pulse type, we examined myosin persistence across different cell rows. Consistently with previous measurements of contractile pulses in the middle of the ventral furrow (Xie and Martin, 2015), pulses close to the ventral midline exhibit persistent myosin; the myosin level after the pulse is higher than the initial baseline (Fig. 6 B, cell row 1). Myosin persistence is associated with a sustained decrease in apical area (Fig. 6 C, cell row 1) (Xie and Martin, 2015). In contrast, myosin pulses at the margin of the mesoderm do not exhibit strong myosin persistence (Fig. 6 B, cell row 7). These myosin pulses accompany cell stretching and do not robustly result in cell apex constriction (Fig. 6 C, cell row 7). Comparing pulse behavior across different ventral-lateral positions, we find a graded decrease in myosin persistence and area stabilization after pulses with distance from the ventral midline (Fig. 6 D).

To further differentiate myosin pulse behaviors, we compared the cross-correlation between the constriction rate and the rate of myosin change in cells along the ventral-lateral axis. Cells closest to the ventral midline exhibit the strongest positive correlation, indicating a correspondence between myosin increase and constriction (Fig. 6 E). Peak correlation is highest in cells along the ventral midline and decreases gradually as distance from the ventral midline increases (Fig. 6 E). Cells ∼7-8 rows from the midline exhibit a small negative correlation, indicating that these cells are prone to not constricting, or may even increase their area during myosin pulses (Fig. 6 F). This behavior is specific to mesoderm cells at this stage because ectoderm cells do not exhibit either a clear positive or negative correlation (Fig. 6 G, cell rows 11, 12). This suggests that the RhoA gradient creates a gradient in cell behaviors coincident with myosin pulses, which we speculate contributes to the transition from constricting to stretching with distance from the midline.

### Actomyosin gradient width regulates furrow curvature and lumen size

To examine the role of the wild-type contractile pattern in successful ventral furrow formation, we tested how disrupting this pattern affects tissue shape. In wild-type embryos with graded constriction, the ventral furrow is a sharp, v-shaped fold with high curvature at its center (Fig. 7A). Previous work showed that globally changing cell fate by altering the *dorsal* gradient to expand the mesoderm results in a flattened depression (Heer et al., 2017). Here, we investigated the role of the actomyosin gradient in tissue folding, by directly modulating RhoA activity without changing mesoderm width. Upon identifying C-GAP, we found that some C-GAP-RNAi embryos have a more C-shaped, less sharp fold (Mason et al., 2016). However, the cause of this abnormal shape was not clear. Therefore, we tested whether widening the gradient by C-GAP-RNAi or RhoGEF2 O/E affects ventral furrow curvature. C-GAP-RNAi embryos in which the zone of uniform constriction is wider have lower central curvature than in wild type (Fig. 7 A and C, Supplemental Fig. 7 A and B). RhoGEF2 O/E embryos also have lower central curvature, suggesting that increased actomyosin gradient width decreases tissue curvature (Fig. 7 B and C, Supplemental Fig. 6 A and B). Indeed, we found that there is an anticorrelation between gradient width and tissue curvature across embryos of different genotypes (Fig. 7 E). Most embryos still fold successfully, although some extreme cases do not manage to internalize the mesoderm. Successful folding in C-GAP-RNAi and RhoGEF2 O/E embryos is associated with a significantly enlarged lumen when invaginated mesoderm forms a tube (Fig. 7D).

**Figure 7:**
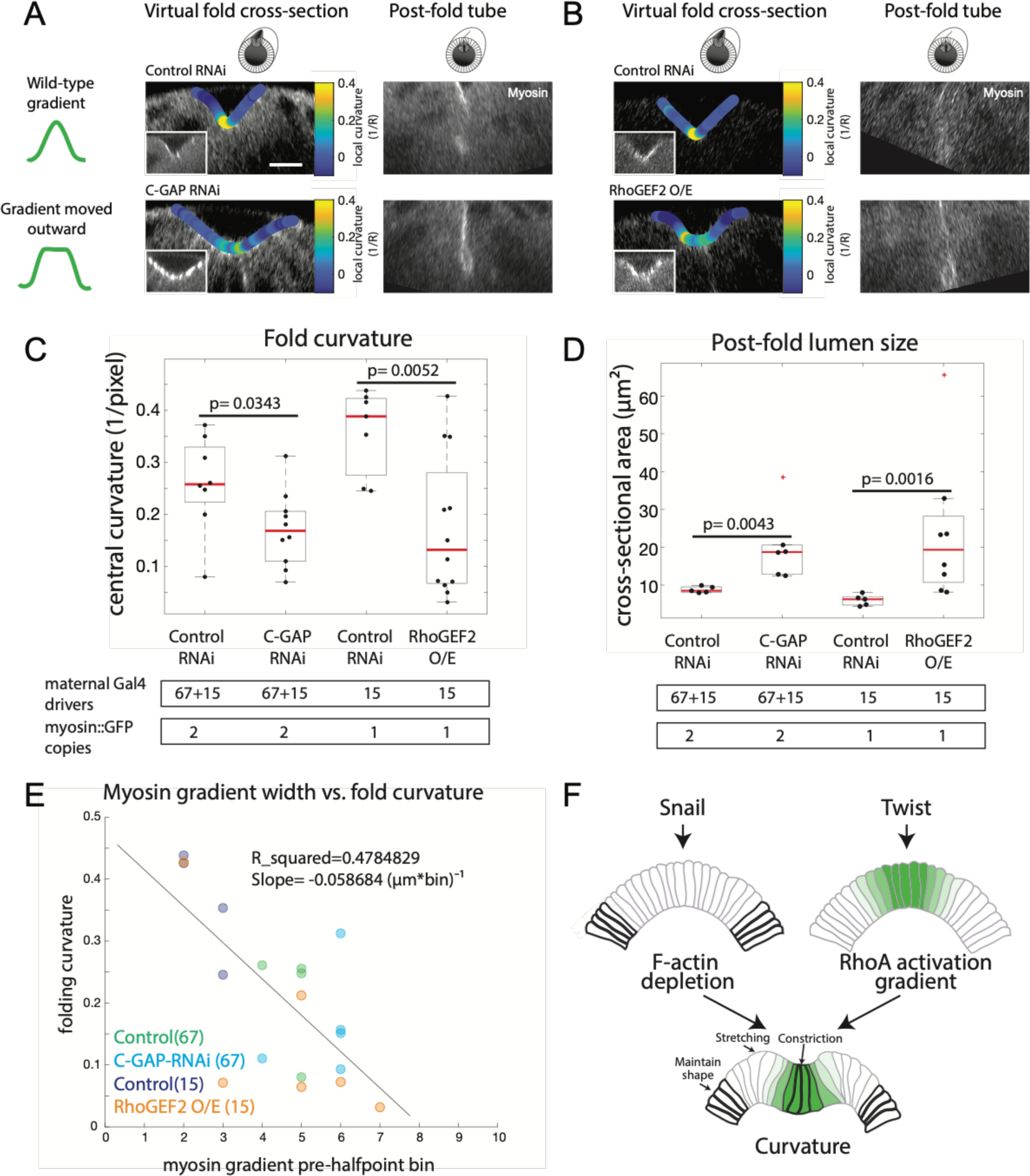
The contractile gradient width affects furrow curvature and post-fold shape. **(A** and **B)** Cross-sectional reslices during and after folding of control (Rh3-RNAi) and C-GAP-RNAi **(A)** or RhoGEF2 O/E **(B)** embryos expressing sqh::GFP (myosin) and gap43::mCherry (membranes). In control embryos, the ventral-lateral cross-section shows a narrow, v-shaped fold with high curvature at the center (local curvature is color-coded on the surface of the fold). The cross-sectional view of the same embryo at a later timepoint shows a tube with a very small lumen. C-GAP-RNAi and RhoGEF2 O/E embryos, which have a widened gradient, display lower central fold curvature and an enlarged tube lumen. Scale bars = 10 µm. Images were rotated to orient ventral side up and black pixels added at corners. **(C-D)** Quantification of curvature at the center of the fold (measured by fitting a circle, three measurements averaged per embryo, C) and lumen size (measured by fitting an ellipse, D) for C-GAP-RNAi and RhoGEF2 O/E embryos with respective controls. Data is represented by box-and-whisker plots overlaid with data points representing each quantified embryo. Bottom and top sides of the box represent 25^th^ and 75^th^ percentile of embryos, respectively. Red midline is the median. P-values are based on pairwise comparison with Mann-Whitney U test. **(E)** Regression analysis of the relationship between myosin gradient width and curvature for control, C-GAP-RNAi and RhoGEF2 O/E embryos. Gradient width was determined as the most lateral bin with mean intensity higher than half-maximal. Curvature was measured as in (C). **(F)** Model for the regulation of tissue-wide patterning in the ventral furrow. Uniform *snail* expression causes uniform F-actin depletion in the mesoderm. Overlapping graded expression of *twist* targets acts via RhoA activation to generate an actomyosin gradient. Fold curvature is driven by the combination of midline contractility and marginal stretching.

To further determine the relationship between junctional F-actin density and curvature, we also decreased RhoA activation. Decreasing RhoA activation by RhoGEF2-RNAi in Utrophin::GFP expressing embryos enabled us to observe the effect of lower F-actin levels in marginal mesoderm cells on folding (Fig. 5 F). These embryos have increased central curvature compared to control embryos (Supplemental Fig. 6 A, B). Although RhoGEF2-RNAi embryos fold successfully, some subsequently exhibit divisions at the embryo surface, suggesting that the mesoderm, which comprises mitotic domain 10, is not completely internalized (Supplemental Movie S3, 4). Overall, our data suggest that the multicellular gradient shape and the F-actin levels in marginal mesoderm cells influence tissue curvature during ventral furrow formation.

## Discussion

Here, we discovered a distinct pattern of junctional F-actin across the *Drosophila* mesoderm and showed how it emerges from the combination of overlapping patterns of transcriptional activity (Fig. 7 F). We showed that Snail-dependent uniform depletion of junctional F-actin throughout the mesoderm, plus Twist-dependent junctional F-actin accumulation in a gradient around the ventral midline, pattern F-actin across the ventral side of the embryo. In addition, we showed that RhoA regulation by the balance of RhoGEF2 to C-GAP determines the actomyosin gradient width. Importantly, the levels and dynamics of F-actin and myosin in distinct cell groups are correlated with their shape changes, leading us to speculate that differences in F-actin density, turnover, and/or myosin persistence between cell groups determine apical constriction vs. stretching behavior across the ventral domain.

### Combination of Twist and Snail creates distinct zones of junctional F-actin density

Prior to folding, we observed through live imaging that Snail expression results in uniform junctional F-actin depletion of mesoderm cells. During folding, Twist expression results in a gradient of junctional F-actin accumulation in a manner that depends on RhoA signaling and adherens junctions. Cells with high F-actin levels tend to maintain their shape or constrict, whereas low-F-actin cells stretch. It was possible that cell strain/stretching or constriction contributed to differences in F-actin density (Latorre et al., 2018). However, the fact that F-actin depletion preludes stretching and is not disrupted when cells are prevented from constricting and stretching their neighbors suggests that F-actin depletion is not a consequence of cell shape. The role of F-actin cortex density in a cell’s response to mechanical stress is well documented (Stricker et al., 2010). In the marginal mesoderm, lower F-actin density in stretching cells is compounded by lower levels of zonula adherens proteins (Dawes-Hoang et al., 2005; Kolsch et al., 2007; Weng and Wieschaus, 2016), both of which could promote the ability of these cells to remodel and stretch in response to stress. In contrast, the neighboring ectoderm and the medial ventral cells have high F-actin and adherens junction density, which may help those cells maintain their shape under stress (Rauzi et al., 2015).

### RhoA activity level determines the width of the actomyosin gradient

Nested within this zone of Snail-mediated F-actin depletion, Twist activity causes actomyosin accumulation via graded RhoA activation. The combination of these overlapping transcriptional patterns allows for domains of different cell behaviors and mechanical properties within a tissue of uniform cell fate (the mesoderm). RhoGTPases are regulated by complex interactions between their (activating) GEFs and (inhibiting) GAPs in many contexts (Denk-Lobnig and Martin, 2019). In the ventral furrow specifically, interactions between C-GAP and RhoGEF2 tune sub-cellular localization and dynamics of the contractile apparatus during folding (Mason et al., 2016). Here, we found that changes to C-GAP or RhoGEF2 levels, at a dose that does not strongly disrupt cell-level organization, change the multicellular pattern of actomyosin levels across the ventral domain. For example, C-GAP-RNAi and RhoGEF2 O/E both widen the myosin gradient and elevate junctional F-actin levels in the marginal mesoderm. Conversely, RhoGEF2-RNAi causes reduced F-actin density in marginal mesoderm cells.

These specific pattern changes can be explained by a model in which there is a graded activator (RhoGEF2) and a uniform inhibitor (C-GAP). In this case, the inhibitor can buffer activation and create a threshold for activation, which would regulate the width of the actomyosin gradient. A prediction of this model is that the gradient widens if overall activator levels increase or inhibitor levels decrease. Furthermore, the upstream activator (RhoGEF2) should be graded over a wider region in wild-type than the downstream myosin. Our measurements of RhoGEF2::GFP show a gradient more similar to the width of apical myosin than upstream pathway components, such as T48 protein (Heer et al., 2017). This could be due to the small signal-to-noise ratio of the RhoGEF2::GFP probe, or due to possible feedback in the RhoA pathway (Priya et al., 2015). In contrast to RhoGEF2 overexpression, RhoGEF2 depletion does not change the myosin gradient width, but results in lower F-actin levels in the marginal mesoderm cells, which could reflect a narrower gradient of F-actin.

Our data show that the gradient shape is tuned by activator-inhibitor balance at the level of direct RhoA regulation. There are other points in the pathway where balance between inhibition and activation is important and could contribute to tissue-wide patterning. In particular, GPRK2, an inhibitor of GPCR signaling, affects myosin organization and cell behaviors (Fuse et al., 2013; Jha et al., 2018). In GPRK2 mutant embryos, apical constriction is expanded, such that lateral mesoderm cells that normally stretch accumulate myosin and constrict (Fuse et al., 2013). This is consistent with a potential role for GPRK2 restricting the contractile gradient.

Further downstream, myosin activity is directly regulated by the balance of ROCK and myosin phosphatase (Munjal et al., 2015; Vasquez et al., 2014). Regulation of tissue-wide properties by overlapping patterns of activator and inhibitors, such as F-actin regulation by Snail and Twist and actomyosin gradient regulation by C-GAP and RhoGEF2, is an intriguing method to create complex spatial patterns of mechanical cell behavior and morphogenesis.

### Fold curvature is tuned by tissue-wide actomyosin patterning

Our disruptions of the actomyosin gradient and the resulting changes in tissue shape suggest that the tissue-wide pattern of the actin cytoskeleton regulates shape. We showed that changing the actomyosin gradient width by modulating the levels of RhoGEF2 or C-GAP effects a change in tissue curvature. Elevating RhoA activity creates a wider gradient and results in C-shaped furrows with low curvature. We showed that the width of the myosin gradient for an individual embryo during flattening is predictive of its furrow and post-fold shape. This is consistent with theoretical work that suggested a broadening contractility would lower tissue curvature (Heer et al., 2017). Another supporting example is that in GPRK2 mutants, all cells within the mesoderm (∼18 cells) constrict and create a U-shaped furrow that often fails to close (Fuse et al., 2013).

That increased contractility in the tissue decreases fold curvature may seem counter-intuitive, but one has to consider the force balance between contractile cells in the tissue. For example, expanding the domain of contractile cells prevents efficient apical constriction at the ventral midline (Chanet et al., 2017; Heer et al., 2017), which we also observed for RhoGEF2 O/E embryos. In many other cases, it has been shown that the successful constriction and invagination of cells depends on neighboring tissue mechanics (Ko et al., 2020; Perez-Mockus et al., 2017; Sui et al., 2018). RhoGEF2-RNAi embryos do not alter the width of the myosin gradient, but have lower junctional F-actin density in marginal mesoderm cells. RhoGEF2 embryos with lower F-actin in the marginal mesoderm have higher furrow curvature. Therefore, it is plausible that the stretching marginal mesoderm cells can promote efficient apical constriction and the creation of a sharp fold at the ventral midline. Together with these past studies, our work emphasizes the importance of understanding tissue-wide patterns in cytoskeletal proteins during folding.

Development generates a multitude of different curvatures and shapes for different contexts. We showed that tissue curvature is sensitive to changes in the pattern of actomyosin within the mesoderm, suggesting that gene expression patterning is an effective way to tune curvature. Given the importance of this patterning mechanism in regulating tissue shape, it is likely that mechanical cell properties are patterned and tuned across the tissue similarly in other developmental contexts with different curvature requirements.

## Materials and Methods

### Fly stocks and crosses

See Supplemental Table S1.

A C-terminal GFP tag was inserted at the endogenous C-GAP locus using CRISPR-Cas9 as previously described (Gratz et al., 2015). Coding sequences for two 15 base pair (bp) gRNAs targeting neighboring sites 5’ of the *rhoGAP71E* gene start codon were cloned into the pU6-BbsI plasmid using the CRISPR Optimal Target Finder (Gratz et al., 2014; Iseli et al., 2007). The donor template plasmid for homology directed repair was generated using Exponential Megapriming PCR (Ulrich et al., 2012). A plasmid backbone (from pHD scarless DS Red) containing an ampicillin resistance gene and an origin of replication was combined with two homology arms (1219 bp and 1119bp, respectively) homologous to the region around the *rhoGAP71E* gene start codon, flanking a *GFP* encoding DNA sequence (kindly provided by Iain Cheeseman) with a N-terminal 4 amino acid-encoding linker region (Ser-Gly-Gly-Ser). Both plasmids were injected into nanos>Cas9 expressing embryos. Surviving adults were crossed to y, w; +; + flies and then screened for mosaic GFP insertion by PCR. Progeny of GFP-positive injected flies were crossed to y, w; +; Dr/TM3 flies and then screened by PCR for the GFP insertion. Successful insertions were further analyzed by sequencing. The fly stock established from their offspring was later back-crossed once to OreR flies in order to eliminate potential off-target mutations.

### Live and fixed imaging

Embryos were collected in plastic cups covered with apple-juice plates. Flies were allowed to lay eggs for 2–4 h at 18-25 °C. The plate was removed and the embryos immersed in Halocarbon 27 oil for staging. Cellular blastoderm stage embryos were collected and prepared for imaging. Embryos were dechorionated with 50% bleach, rinsed with water, and then mounted on a slide with embryo glue (Scotch tape resuspended in heptane), with the ventral side facing upwards. A chamber was made with two no. 1.5 coverslips as spacers, a no. 1.0 coverslip placed on top, and the chamber was filled with Halocarbon 27 oil before imaging. Images were acquired on a Zeiss 710 microscope with an Apochromat 40×/1.2 numerical aperture W Korr M27 objective.

Immuno-and phalloidin staining was performed using standard methods (Martin et al., 2009). Embryos were fixed with 4% paraformaldehyde/heptane for 30 min, devitellinized manually, stained with phalloidin, primary antibodies and appropriate fluorescently tagged secondary antibodies, and mounted in AquaPolymount (Polysciences, Inc.). Anti-snail (rabbit, 1:100, M. Biggin, Lawrence Berkeley National Lab), anti-GFP (rabbit, 1:500; Abcam, ab290) and anti-E-cadherin (rat, 1:50; DHSB) antibodies and AlexaFluor568 phalloidin (Invitrogen) were used. All imaging was carried out on a Zeiss 710 confocal microscope with a Plan-Apochromat 40×/1.2 numerical aperture W Korr M27 objective.

For imaging settings, refer to Supplemental Table S2.

### Gradient analysis

Analysis was done as described in Heer et al. 2017. All image analysis was performed in Fiji (http://fiji.sc) (Schindelin et al., 2012) and MATLAB (MathWorks). Custom software for image processing is available upon request.

#### Definition of developmental timing

Wild-type embryos were staged based on the time of folding. The accuracy of this method was confirmed by comparing constricted areas per bin at the selected time point. For embryos with disrupted constriction and folding, an analogous time point was chosen relative to the beginning of myosin/ fluorescence accumulation.

#### Shell projection and thresholding to measure apical fluorescence intensity

Shell projections of the apical surface were made to capture the embryo surface. First, cytoplasmic background signal (defined as the mean cytoplasmic signal plus 2.5 standard deviations) was subtracted from the myosin channel (Martin et al., 2009; Vasquez et al., 2014). For non-myosin fluorescent signal (Fig. 3), the cytoplasmic background subtraction was adjusted to account for differences in signal-to-noise ratio for different fluorescent markers (RhoGEF2-GFP: mean + 2 standard deviations (SDs); aniRBD-GFP: 2 SDs, rok-GFP: 2.5 to 3 SDs).

The maximum myosin (or other apically enriched fluorescent) signal intensity in the *z*-plane was used to generate a rough map of the embryo surface. A Fourier transform was used to generate a smooth continuous surface. Myosin signal was averaged over the 4 µm above the surface of detected maximum intensity and membrane signal was the sum of the signal from 1 µm below the surface. A gaussian blur filter (radius 1 pixel for fluorescent signal, 0.7-1 membranes) was applied after shell projection to reduce noise.

Shell projections from live and immunostained images were then segmented using an existing MATLAB package, Embryo Development Geometry Explorer (EDGE) (Gelbart et al., 2012). Membrane signal (Gap43::mCherry) or cortical actin (phalloidin) projections were used to detect cell boundaries (and track cells in time for live images). Errors in segmentation were corrected manually. Our segmentation algorithm was used to determine centroid position, cell diameter, cell area, cell perimeter of segmented cells as well as total myosin/fluorescent signal per cell based on the corresponding myosin/fluorescent channel projection.

#### Defining cell bins

For all image quantifications, data was aggregated into ‘cell bins’ (Heer et al., 2017). Cells were assigned to bins based on the ventral-lateral position of the cell centroid relative to the ventral midline. The ventral midline (VM) was defined as the position at which the furrow closes. In fixed images or for embryos that did not fold (or rotated while folding), the position of the VM was determined by symmetry of the fluorescent signal. Live images with several segmented time points were binned based on initial position of the cell centroid before constriction and folding; the boundaries of the bins were set based on the average cell diameter along the ventral-lateral axis. For images in which cells had already started to constrict, the width of each bin was set manually (but still relative to average cell diameter) to approximate the width of cells at that ventral-lateral position. We used MATLAB to generate box-and-whiskers plots depicting the distribution of data, overlaid with the mean of each bin. For boxplots, bottom and top sides of the box represent 25th and 75th percentile of cells, respectively. Outliers were defined as values 1.5 times bigger than the interquartile range. Fluorescent signal was normalized by dividing by the mean of the bin with highest average intensity, to adjust for variability in imaging conditions.

#### Junctional F-actin quantification

For apical projections of phalloidin staining (Fig.1 A, 2 A and 4 B), embryos were shell-projected and cells tracked similarly to myosin and other markers (see above), but no background subtraction was required because of the low background staining for phalloidin stained embryos. For analysis, junctional signal was extracted from the apical shell projection by only using signal within 2 µm of the tracked cell boundaries. This total junctional F-actin signal was divided by the cell perimeter to obtain junctional F-actin density independently of cell circumference.

Because mesodermal F-actin depletion was most obvious subapically, not apically, with fluorescently tagged Utrophin markers in live embryos (Fig. 2 B, 5 C and 7 C), those movies were “shell-projected” in Fiji by generating a Z-reslice (1 µm per slice) and then doing a second reslice along a manually drawn, subapical segmented line that followed the ventral-lateral curvature of the embryo surface (Supplemental Fig. 2). This allowed us to account for specific ventral-lateral curvature and get a subapical projection at a consistent apical-basal depth. The center part along the AP axis of the shell-projection, where anterior-posterior (AP) curvature is small, was used to quantify F-actin (Utrophin::mCherry or ::GFP) intensity along the ventral-lateral axis. For analysis of the tissue-wide pattern, junctional signal was extracted from the subapical shell projection by only using signal within 2 µm of the tracked cell boundaries. This total junctional F-actin signal was divided by the cell perimeter to obtain junctional F-actin density independently of cell size.

### Pulsing analysis

Images of live embryos with myosin and membrane markers during folding were obtained as above, but with faster scan speed and smaller z-depth, to obtain time steps between 6 and 10 seconds, which is sufficient to capture typical myosin and area pulsing behavior (Martin et al 2009). Cells across the mesoderm were tracked over time using EDGE and cell area and myosin intensity were exported. In Matlab 2019a, we detected peaks within individual cells of maximal myosin intensity increase by smoothing with a moving average filter and then detecting local maxima. The myosin intensity and cell area behavior about 100 seconds before and after each maximum was saved as a trace. Pulse traces were averaged for each cell bin to identify average behavior based on ventral-lateral cell position. Myosin persistence was defined as the minimum myosin intensity within each trace after a myosin pulse minus the minimum myosin intensity before the same pulse. Persistence of area constriction was defined as the maximum cell area after a myosin pulse minus maximum cell area before the same pulse. Persistence values were averaged by bin and plotted in Matlab for three individual embryos.

To analyze how the relationship between area and myosin behaviors changes based on cell position, we cross-correlated myosin and area behavior (as described in Martin et al. 2009). We used the xcorr function in Matlab to cross-correlate the change in myosin intensity (myosin intensity at a timepoint minus myosin intensity at the previous time point) with constriction change (cell area at a timepoint minus cell area at the next time point, i.e. positive value if constricting, negative if stretching) for each cell trace. Cross-correlations for each cell were averaged by bin (distance from the midline) and plotted.

### Curvature analysis

Movies with at least 20 µm z-depth were resliced in FIJI to create transverse virtual cross sections (1 µm thickness) of the ventral furrow all along the embryo’s AP axis. Apically enriched myosin signal was used to trace the apical surface of the folding tissue with the freehand line tool in Fiji. Traces were made for a cross section at the center of the AP axis of the embryo as well as for cross-sections 20 µm anterior and posterior, respectively. The timepoint of measurement was chosen based on invagination depth (about 10 µm). The XY coordinates of each trace were imported into Matlab R2019a. After manually determining the position of the ventral midline, a circle fit (Nikolai Chernov (2020). Circle Fit (Taubin method) (https://www.mathworks.com/matlabcentral/fileexchange/22678-circle-fit-taubin-method), MATLAB Central File Exchange. Retrieved February 5, 2020) was applied to the central part of the trace, 2.5 µm left and right of the midline. The inverse of the fitted circle radius was defined as the fold curvature. The three traces taken from each embryo were averaged to obtain the fold curvature value for that embryo. In addition, local curvature was visualized using LineCurvature2D (Dirk-Jan Kroon (2020). 2D Line Curvature and Normals (https://www.mathworks.com/matlabcentral/fileexchange/32696-2d-line-curvature-and-normals), MATLAB Central File Exchange. Retrieved March 16, 2020.) in Matlab.

Post-folding lumina were visualized from central cross-sections of the same embryos whose curvature had been analyzed and measured for a single AP position. Lumen area was determined by manually fitting an ellipse to the lumen in each cross-section in Fiji and measuring its area.

Curvature and lumen measurements were compared between genotypes using the non-parametric two-sample Mann-Whitney U test (‘ranksum’ command in Matlab).

### Regression analysis

All linear regression fits (Fig. 1 E, 7 D) were performed in Matlab R2019a using the fitlm command. The original data as well as the best fit line were plotted and the R-squared value was reported as a measure of fit.

For myosin gradients, the position of the most lateral bin with myosin levels above half-maximal was used to describe gradient width. This bin position was then compared to central curvature measurements of the same embryo at a later timepoint (at ∼10 µm invagination depth, analyzed with circlefit).

## Acknowledgments

We thank N. Perrimon, L. Perkins and the Transgenic RNAi Project at Harvard Medical School (National Institutes of Health/National Institutes of General Medical Sciences R01-GM084947) for providing transgenic RNAi stocks. We thank V. Tserunyan for the software adapted here to analyze myosin pulses. We thank B. Adhikary for help with analysis of F-actin levels. We thank current and former Martin lab members for helpful discussions and feedback. This work was supported by a grant from the National Institutes of General Medical Sciences: R01GM125846 to A.C.M.

---

**Supplemental Figure 1.**
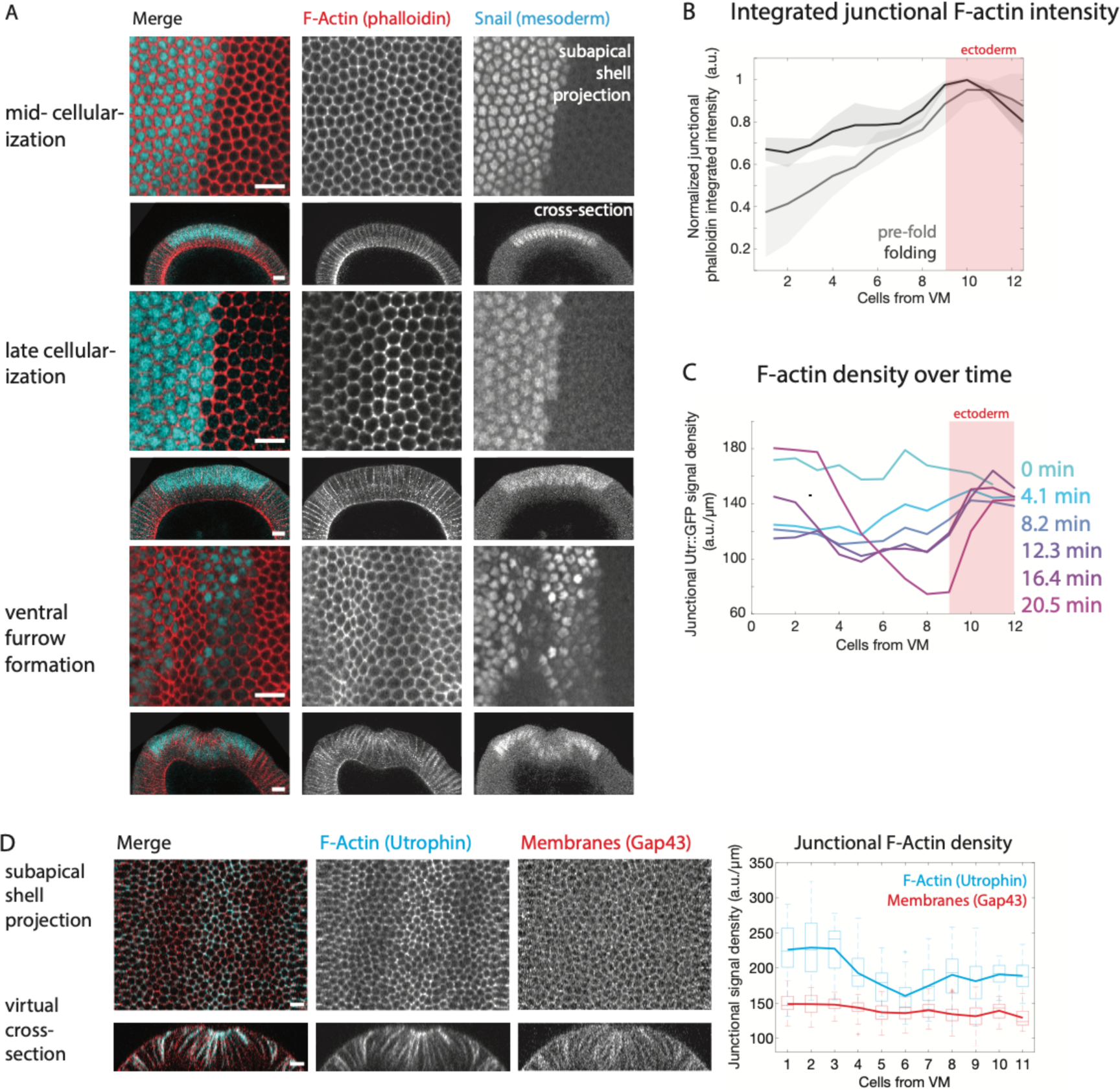
F-actin is depleted in the mesoderm during cellularization. **(A)** F-actin depletion in the mesoderm appears towards the end of cellularization. Images are subapical shell projections (top) and cross-sections (bottom) of wild-type embryos at different stages stained with phalloidin (red) and anti-snail antibody (cyan). Scale bars = 10 µm. Cross-sectional images were rotated to orient ventral side up and black pixels added at corners. Brightness and contrast were adjusted individually to best display the intensity range in each image. **(B)** Integrated junctional F-actin intensity per cell before and after folding (mean and standard deviation, for same embryos as Figure 1 D). Mean junctional F-actin intensity was calculated by segmenting cells and integrating F-actin intensity around the cell periphery. All traces were normalized to their highest-mean cell bin before averaging. **(C)** Mean subapical F-actin junctional density by cell row over time for a single embryo marked with Utrophin::GFP, from mid-cellularization (0 min) to during folding (20.5 min). F-actin density stays mostly constant in the ectoderm, but decreases in the mesoderm during cellularization and then increases in a gradient around the ventral midline. **(D)** Left: F-actin marked by the fluorescently marked Utrophin::GFP (cyan) is depleted in the marginal mesoderm, but a general membrane marker, Gap43::mCherry (red), is not. Images show both markers in subapical shell projections (gaussian blur radius=1) and cross-sections. Scale bar = 10 µm. Right: Quantification by bin of subapical Utrophin::GFP (top) and Gap43::mCherry (bottom) junctional fluorescent intensity, normalized by perimeter. Data is represented by box-and-whisker plots where each bin is a cell row at a given distance from the ventral midline (at least 21 cells per row analyzed, median 38 cells). Bottom and top sides of the box represent 25^th^ and 75^th^ percentile junctional intensity per cell, respectively. Midline is the median and red points are outliers.

**Supplemental Figure 2.**
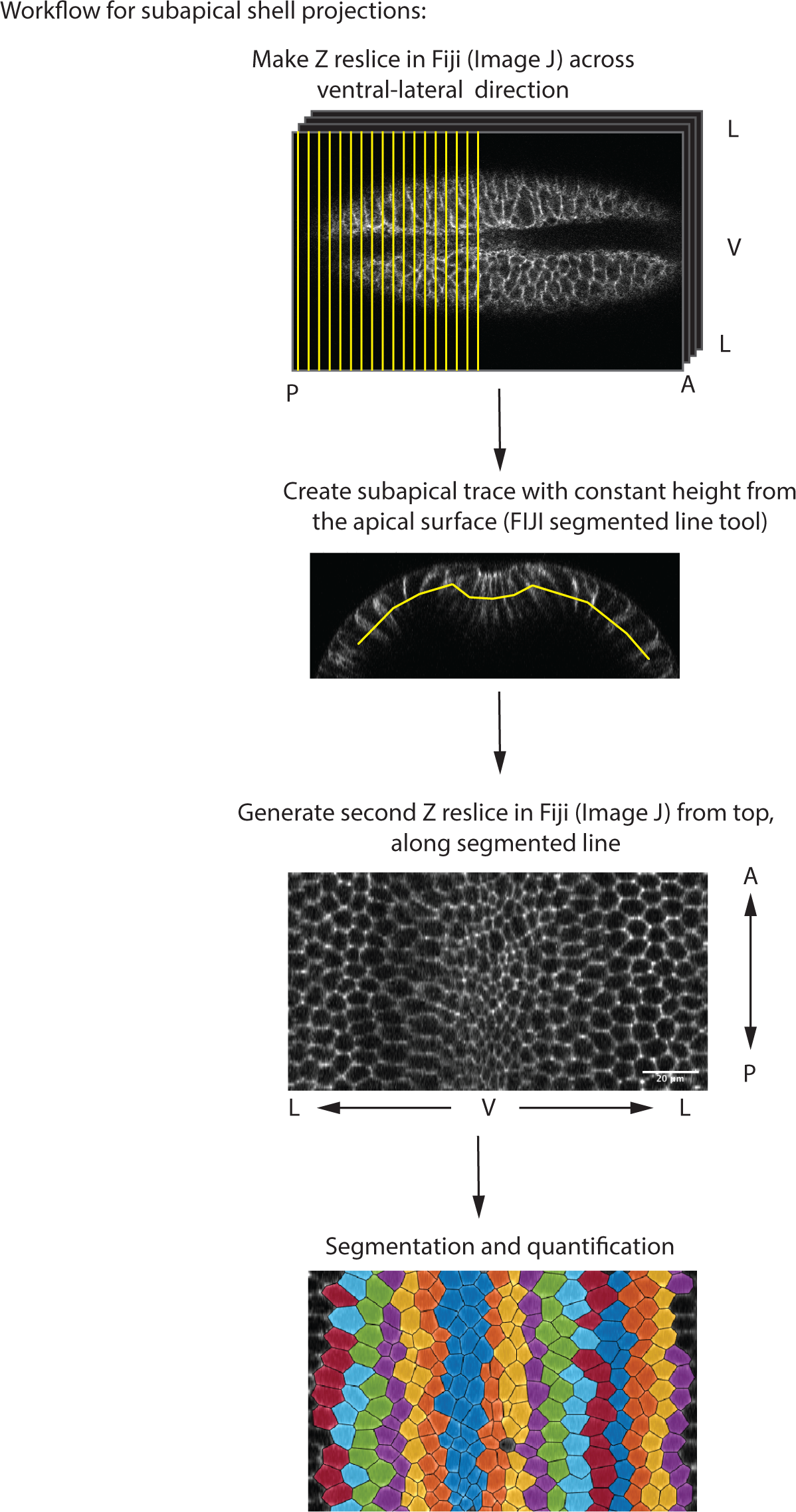
Workflow for subapical shell projections in FIJI. Confocal image stacks are resliced along the ventral-lateral axis to get a cross-sectional view. A subapical trace is then drawn manually and the embryo is resliced a second time along the trace. The central part of the embryo along the AP axis is then used to segment cells and quantify intensity along the ventral-lateral axis.

**Supplemental Figure 3:**
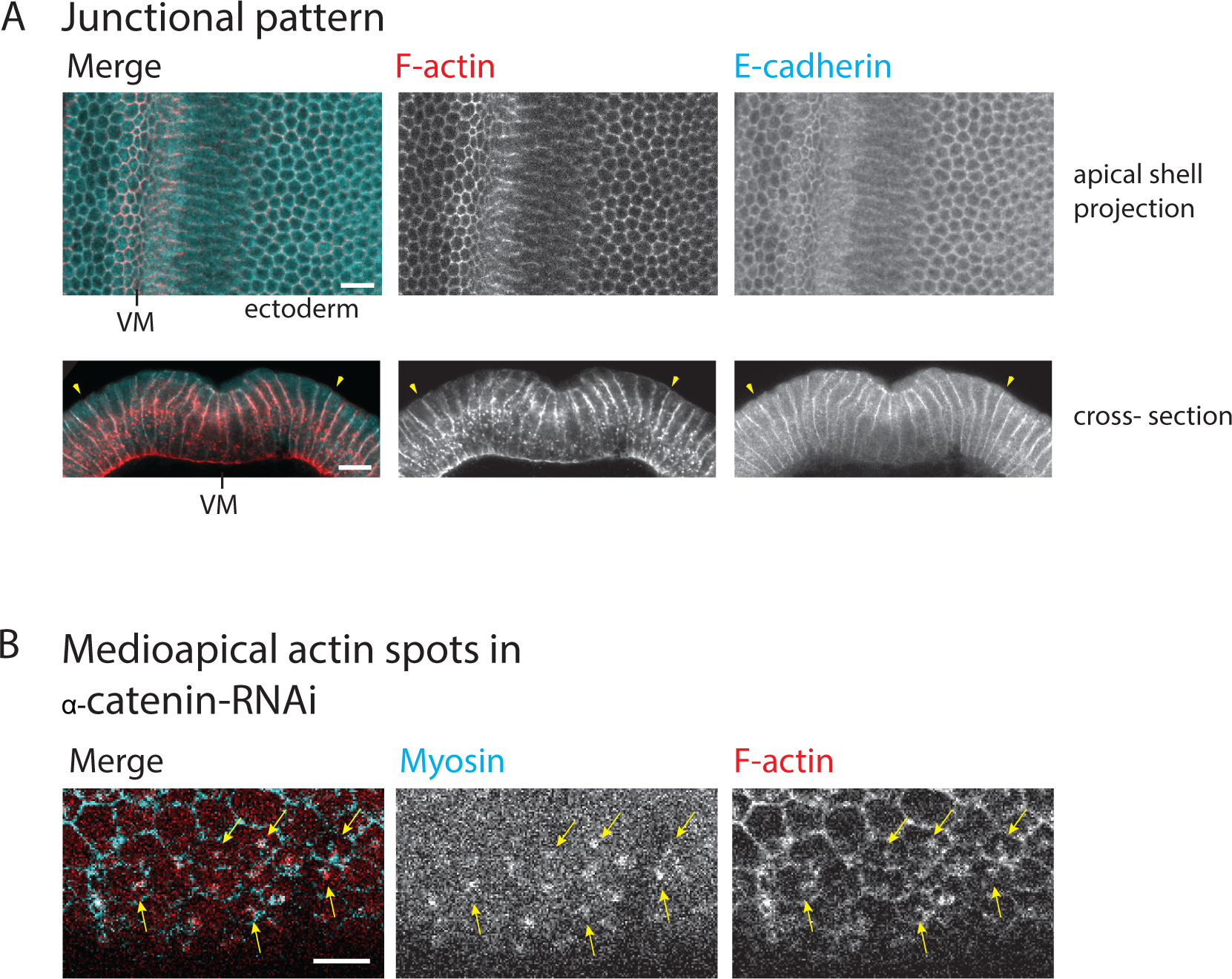
F-actin is colocalized with junctions in wild-type and with medioapical myosin spots in α -catenin-RNAi. **(A)** Top: apical shell projection images of fixed control RNAi embryo stained with phalloidin (red) and anti-E-cadherin antibody (cyan). Bottom: Cross-section images of fixed wild-type embryo stained with phalloidin (red) and anti-E-cadherin antibody (cyan). Yellow arrows mark mesoderm-ectoderm border. Scale bars = 10 µm. Cross-sectional images were rotated to orient ventral side up and black pixels added at corners. **(B)** Images of ventral apical surface of fixed embryo expressing sqh::GFP (Myosin light chain marker), stained with phalloidin (red) and anti-GFP (cyan) antibody (same embryo as in Figure 3 D). Yellow arrows indicate medioapical spots in which myosin and F-actin are colocalized. Scale bar = 10 µm.

**Supplemental Figure 4.**
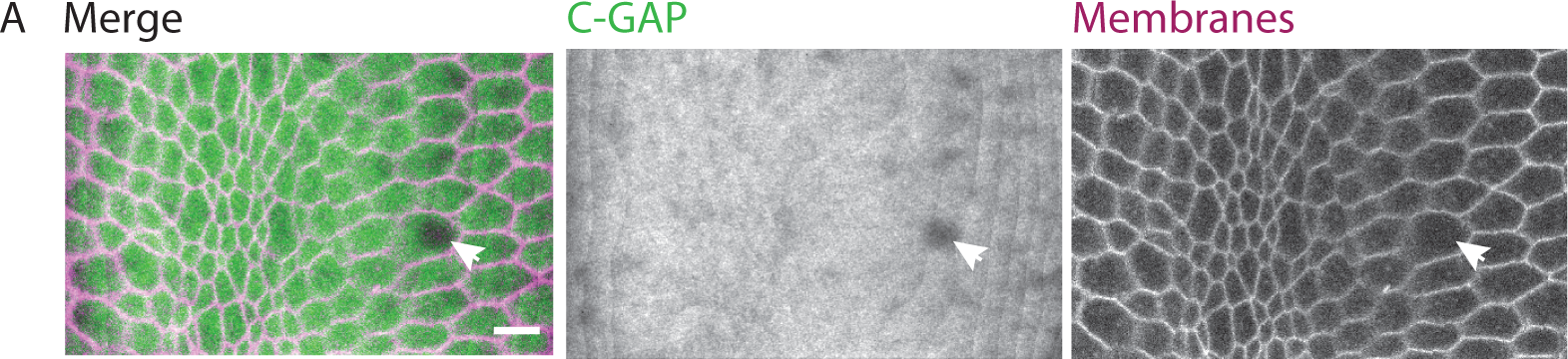
C-GAP is uniformly cytoplasmic across the ventral furrow. **(A)** Apical shell projection of an embryo expressing GFP::C-GAP (green) and gap43::mCherry (membranes, magenta). White arrows indicate apical nucleus with no GFP::C-GAP signal. Scale bar = 10 µm.

**Supplemental Figure 5:**
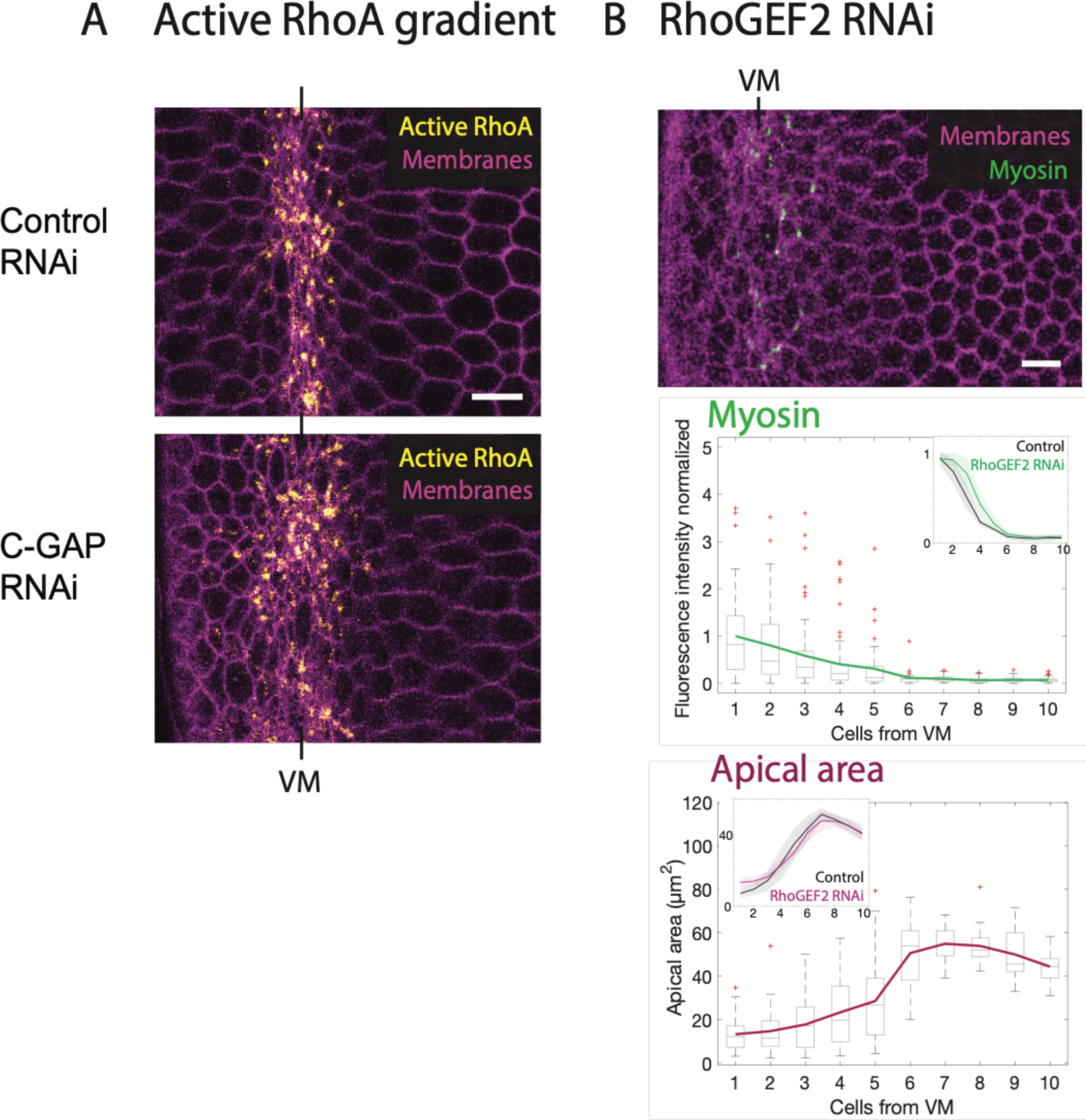
C-GAP-RNAi expands the active RhoA zone and RhoGEF2-RNAi decreases myosin levels, but there is still a gradient. **(A)** Images (apical shell projection) of control (Rh3-RNAi) and C-GAP-RNAi embryos expressing Anillin Rho-binding domain::GFP (active RhoA, yellow) and Gap43::mCherry (Membranes, magenta). Scale bar = 10 µm. **(B)** Top: Image (apical shell projection) of RhoGEF2-RNAi embryo expressing sqh::GFP (Myosin, green) and Gap43::mCherry (Membranes, magenta). Scale bar = 10 µm. Bottom: Quantification of apical area (magenta) and normalized apical, active myosin (green) as a function of distance from ventral midline (at least 23 cells per cell row were analyzed; median 38 cells). Data is represented by box-and-whisker plots where each bin is a cell row at a given distance from the ventral midline. Bottom and top sides of the box represent 25^th^ and 75^th^ percentile of cells, respectively. Midline is the median and red points are outliers. Inset shows average cell behavior (and standard deviation) from 5 different RhoGEF2-RNAi and 4 different control embryos. Short hairpin RNAs were driven by Mat 67 and 15 Gal4 drivers. Control embryos are the same as in Figure 5 D.

**Supplemental Figure 6:**
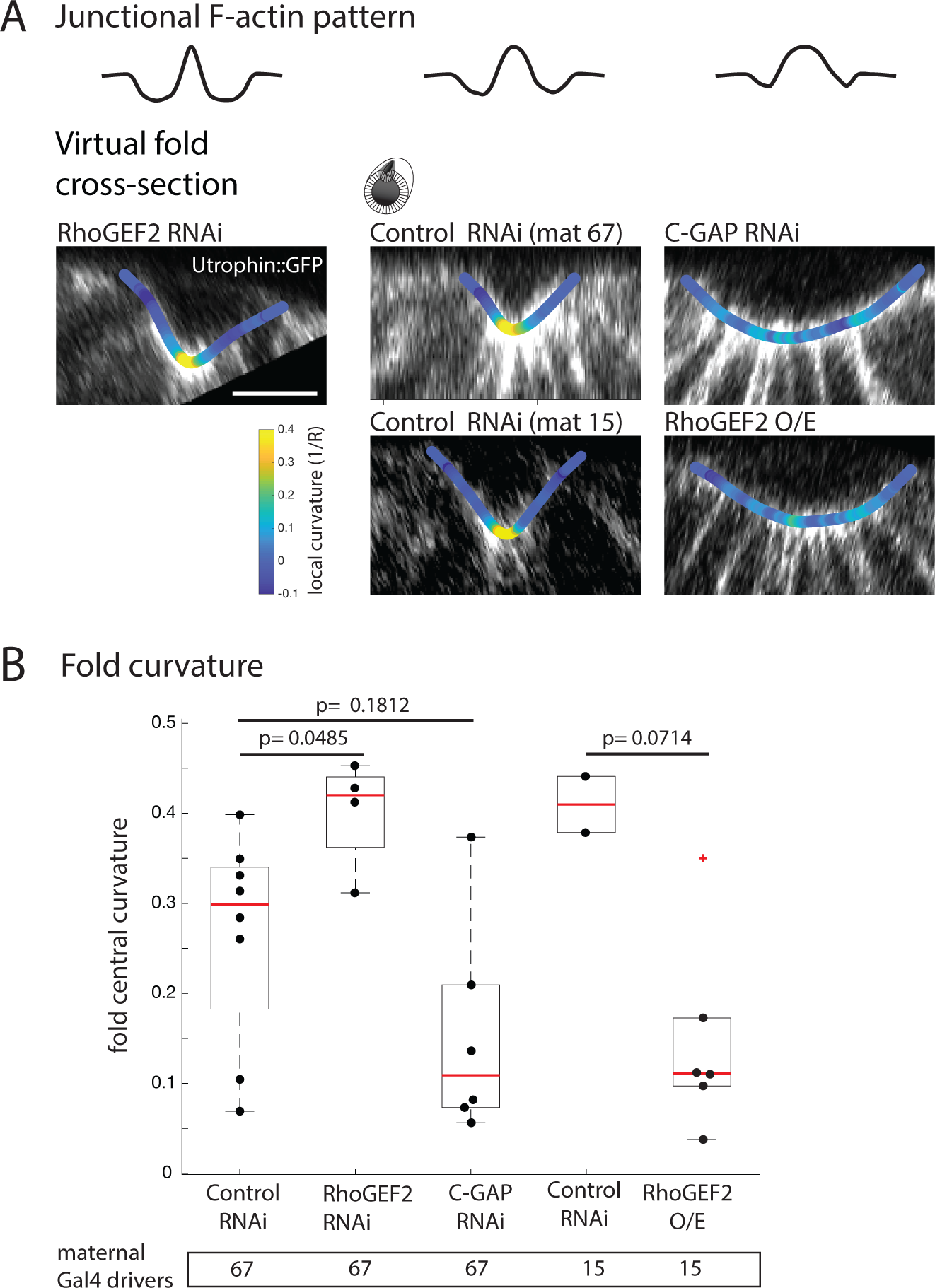
Junctional F-actin levels in the marginal mesoderm affect fold curvature. **(A) Top:** Cross-sections during folding of RhoGEF2-RNAi, control (Rh3-RNAi) and C-GAP-RNAi embryos expressing Utrophin-GFP. In cross-section, local curvature is color-coded on the surface of the fold. Scale bars = 10 µm. Bottom: Cross-sections during folding of control (Rh3-RNAi) and RhoGEF2 O/E embryos expressing Utrophin-GFP. Images were rotated to orient ventral side up and black pixels added at corners. **(B)** Quantification of curvature at the center of the fold (measured by fitting a circle, three measurements averaged per embryo) for control (Rh3-RNAi), RhoGEF2-RNAi, control (Rh3-RNAi) and C-GAP-RNAi, as well as control (Rh3-RNAi) and RhoGEF2 O/E embryos. Maternal GAL4 drivers used to drive short-hairpin RNAs or RhoGEF2 are indicated. Data is represented by box-and-whisker plots overlaid with data points from all quantified embryo. Bottom and top sides of the box represent 25^th^ and 75^th^ percentile, respectively. Midline is the median. P-values are based on pairwise comparisons with Mann-Whitney U test.

**Supplemental Movie S1 (related to Fig. 4)**. Shell-projection (with background subtraction and gaussian blur) of control (Rh3-RNAi) embryo expressing sqh::GFP (myosin) and gap43::mCherry (membranes) during ventral furrow formation.

**Supplemental Movie S2 (related to Fig. 4)**. Shell-projection (with background subtraction and gaussian blur) of α-catenin-RNAi embryo expressing sqh::GFP (myosin) and gap43::mCherry (membranes) trying to initiate ventral furrow formation.

**Supplemental Movie S3 (related to Fig. 7)**. Maximum intensity projection (38 µm depth) of control (Rh3-RNAi) embryo expressing sqh::GFP (myosin) and gap43::mCherry (membranes), during and after ventral furrow formation.

**Supplemental Movie S4 (related to Fig. 7)**. Maximum intensity projection (38 µm depth) of RhoGEF2-RNAi embryo expressing sqh::GFP (myosin) and gap43::mCherry (membranes), during and after ventral furrow formation. Note divisions of mitotic domain 10 visible at the embryo surface after folding.

## Notes

### Competing Interest Statement

The authors have declared no competing interest.

