## Supplemental Table S1 for "Combinatorial patterns of graded RhoA activation and uniform F-actin depletion promote tissue curvature"

| Stock Nr. | Genotype | Source/Reference |
|---|---|---|
| 1 | w; Gap43::mCherry(attp40); Sqh::GFP | Martin et al., 2010 |
| 2 | y, w[67c], sqh[AX3], cv; Sqh::GFP[42] | Bloomington Drosophila Stock Center |
| 3 | OreR | Bloomington Drosophila Stock Center |
| 4 | y,w;+;C-GAP-GFP | This study |
| 5 | y[1] sc[*] v[1]; P{y[+t7.7] v[+t1.8]=TRiP.GL01052}attP2 (Rh3 shRNA control line) | Perkins et al., 2015 |
| 6 | Mat 67, UtrABD-GFP | Jodoin and Martin, 2016 |
| 7 | UtrABD-GFP;+ | Rauzi et al. 2010 |
| 8 | w; gap43::mCherry/TM3, Sb[1] | Bardet et al. 2013 |
| 9 | UtrABD-mCH | Rauzi et al. 2010 |
| 10 | Halo-sna/CyO-sqhGFP | Martin et al., 2009 |
| 11 | halo, twist[ey53] /CyO, Sqh::GFP | Martin et al., 2009 |
| 12 | w; mat67, Sqh::GFP; mat15, Gap43::mCherry/(TM3, Sb[1]) | Vasquez et al., 2014 |
| 13 | sqhGFP; P{y[+t7.7] v[+t1.8]=TRiP.GL01052}attP2 (Rh3 shRNA control line) | This study |
| 14 | Alpha-catenin RNAi (HMS), sqh-GFP | This study (original Alpha-catenin-RNAi stock: Perkins et al., 2015) |
| 15 | w; mat67; mat15 | Vasquez et al., 2014 |
| 16 | y w hs-flp; FRT42BG13 RhoGEF2[1.1]/CyO; GFP::Rhogef2 BAC(VK33) | Mason et al. 2016 |
| 17 | w[1118]; Df(2R)ED2747, P{w[+mW.Scer\FRT.hs3]=3'.RS5+3.3'}ED2747/SM6a; GFP::Rhogef BAC(VK33), Gap43::mCherry | Mason et al. 2016 |
| 18 | Ubi-AniRBD-GFP; gap43mCH | This study (original Ubi>AniRBD-GFP stock: Munjal et al. 2015) |
| 19 | Ubi-RokGFP; gap43mCh | Bardet et al. 2013 |
| 20 | y,w; Gap43::mCherry/Cyo;C-GAP-GFP | This study |
| 21 | sqhGFP; P{TRiP.HMS00412}attP2 (RhoGAP71E, C-GAP shRNA) | This study (original C-GAP-RNAi stock: Perkins et al., 2015) |
| 22 | y[1] w[*]; P{UASp-T7.RhoGEF2}5 (RhoGEF2 overexpression line) | Bloomington Drosophila Stock Center |
| 23 | y,w; Sqh::GFP; mat15, Gap43::mCherry/(TM3, Sb[1]) | Vasquez et al., 2014 |
| 24 | y[1] sc[*] v[1]; P{TRiP.HMS01118}attP2 (*RhoGEF2 shRNA*) | Perkins et al., 2015 |
| 25 | mat15, Utr::mCh | This study |
| 26 | mat67; mat15, gap43::mCh | Vasquez et al., 2014 |
| 27 | Ubi-AniRBD-GFP; 71E HMS | This study |
| 28 | Ubi-AniRBD-GFP; RH3GL | this study |


| Figure | Cross or stock | Comments |
|---|---|---|
| 1A | Stock # 1 x 2 (Virgins x males) | |
| 1B-E | 3 | |
| 1B-E | 4 | |
| S1A,B | 3 | |
| S1C | 5x6 | |
| S1D | 7x8 | |
| 2A | 3 | |
| 2B,C,F | 9x10 | mutants selected by halo phenotype |
| 2D,E,F | 9x11 | mutants selected by halo phenotype |
| 3A-C,E | 12x13 | |
| 3A-C,E | 12x14 | |
| 3D-E | 5x15 | |
| 3D-E | 14x15 | |
| S3A | 13x15 | |
| S3A | 3 | |
| S3B | 14x15 | |
| 4B,C | 16x17 | |
| 4B,C | 3x18 | |
| 4B,C | 3x19 | |
| S4 | 4x20 | |
| 5A,B | 12x13 | |
| 5A,B | 12x21 | |
| 5C,D | 22x23 | |
| 5C,D | 5x23 | |
| 5E,F | 5x6 | |
| 5E,F | 6x21 | |
| 5E,F | 6x24 | |
| 5G,H | 22x25 | |
| 5G,H | 5x25 | |
| S5A | 26x27 | |
| S5A | 26x28 | |
| S5B | 23x24 | |
| S5B | 5x23 | |
| 6 | 1x2 | |
| 7A,C,D | 12x13 | |
| 7A,C,D | 12x21 | |
| 7B,C,D | 22x23 | |
| 7B,C,D | 5x23 | |
| S7 | 5x6 | |
| S7 | 6x21 | |
| S7 | 6x24 | |
| S7 | 22x25 | |
| S7 | 5x25 | |

**F2 embryos imaged from these crosses, using above stock numbers/genotypes. Non-balancer females were used for cages.**
