## Supplemental Table S2 for "Combinatorial patterns of graded RhoA activation and uniform F-actin depletion promote tissue curvature"

**Imaging settings**

| Fluorescent tag combination | Figures | Excitation laser wavelength (nm) | Detector start wavelength (nm) | Detector end wavelength (nm) | Detector Gain (a.u.) | Laser power (%) | Pinhole size ($x10^{-4}$m) |
|---|---|---|---|---|---|---|---|
| sqh::GFP (Myosin) | 1A, 3A-C, S3C, 5A, S5B, 6, 7 | 488 | 493 | 537.57 to 586 | 700 to 874.081 | 2 to 7 | 89.944 |
| gap43mCh (Membranes) | 1A, S1D, S3, 3A-C, S3C, 4, 5A, S5A+B, 6, 7 | 561<br>594 | 569.57 to 603.82<br>600.32 to 614.82 | 696 to 698.26<br>696 | 763 to 866<br>840 to 856.321 | 1.5 to 6.5<br>5 to 7 | 89.944<br>89.944 |
| Utrophin::GFP | S1C,D, 5D-F, S7 | 488 | 493 | 562.07 to 598 | 769.995 to 850 | 2 to 4 | 89.944 |
| Utrophin::mCh | 2B-E | 561 | 578 | 696 | 770-800 | 4.3 to 5 | 89.944 |
| RhoGEF2::GFP | 4B | 488 | 493 | 561 | 850 | 12 | 89.944 |
| Anillin::GFP | 4C, S5A | 488 | 493 | 561 | 850 | 2.5 to 22 | 89.944 |
| Rok-GFP | 4D | 488 | 493 | 561 | 850 | 7 | 89.944 |
| C-GAP-GFP | S4 | 488 | 493 | 565 | 900 | 16 | 89.214 |
| Phalloidin Alexa Fluor 568 | 1B-D, S1A, 3D, S3A, B | 561 | 569 to 574 | 647 to 712 | 600 to 700 | 1 to 2 | 37.175 to 77.853 |
| Alexa Fluor 488 secondary antibody | S1A (snail), 2A (snail) | 488 | 493 | 574 | 500 to 700 | 2 to 4 | 38.114 to 89.213 |
| Alexa Fluor 647 secondary antibody | S3A (E-cadherin) | 633 | 647 | 755 | 800 | 5 | 42.175 |
| sqhGFP fixed | S3B | 488 | 493 | 569 | 800 | 8 | 37.175 |
